# Linking plant growth promoting arbuscular mycorrhizal colonization with bacterial plant sulfur supply

**DOI:** 10.1101/2021.06.22.449381

**Authors:** Jacinta Gahan, Achim Schmalenberger

## Abstract

Sulfur (S) exists in organically bound complexes (∼95%), predominantly as sulfonates, and are not directly plant available. Specific soil bacteria can mobilise sulfonates but very little is known about these bacteria in the hyphosphere. Since mycorrhizal fungi support growth of the majority of land plants, hyphosphere desulfonating bacteria may be of substantial benefit to the plant host. This study analysed the effect of AM inoculation with *Rhizophagus irregularis* (former *G. intraradices,* Glomus) and a mix of six AM species (Mixed) on PGP, microbial communities and sulfonate mobilising bacteria with *L. perenne*, *Agrostis stolonifera* and *Plantago lanceolata* as plant hosts in bi-compartmental microcosms and *A. stolonifera* in PGP pot experiments. AM inoculation significantly increased plant growth, percentage root colonisation and the quantity of cultivable desulfonating bacteria in the hyphosphere over pre-inoculated soil for all plants. Community analysis via PCR-DGGE revealed significantly different bacterial and fungal communities post inoculation. Analysis of the sulfonate mobilising *asfA* gene revealed a significantly altered community and novel bacterial isolates with this important functional ability post-inoculation. The results demonstrate that AM inoculation increased plant biomass yield, AM root colonisation and altered bacterial and fungal community dynamics in the hyphosphere. AM inoculated microcosms had an increased abundance of desulfonating bacteria that may be beneficial for plant-S supply.

**Research highlights:**

- Inoculation with AM fungi was shown to promote plant growth and harbour larger populations of sulfonate mobilising bacteria.
- Post-inoculation hyphospheric bacterial and saprotrophic fungal communities were shown to differ significantly in composition and abundance.
- Analysis of sulfonate mobilising bacteria revealed novel presumptive species in possession of the *asfA* gene associated with AM hyphae.
- AM inoculation was shown to significantly impact the *asfA* positive bacterial community composition.

## 1. Introduction

Agricultural land has been subjected to human induced degradation in recent years as a result of intensification of agricultural practices leading to ecosystem alterations including soil degradation and erosion (Barrios, 2007). Restoration and sustainable management of agricultural land is a necessity given the growing global population, estimated to double by 2050, and increased demand from agricultural land projected to be 60-110% of the current rate (Ray *et al*., 2013).

Arbuscular Mycorrhizal (AM) fungi are ubiquitous soil microorganisms that form endosymbiotic partnerships with most land plants. They exist in almost all terrestrial ecosystems where they may be involved in sustainable ecosystem services, for example, in nutrient depleted agricultural soils (Öpik *et al*., 2006). AM fungi are close to being eradicated from agricultural land under current practices which include; tillage (Johansson *et al*., 2004), crop rotation (Plenchette *et al*., 2005) and application of agrochemicals (Bünemann *et al*., 2006).

In recent years, the use of commercially available AM inocula to restore native AM fungal communities has been investigated with the objective of maximising the many biotic and abiotic benefits of an intact symbiosis. These benefits include i) improved nutrient acquisition, ii) stabilisation of soil aggregates, iii) prevention of erosion and iv) alleviation of potential plant stress factors (Wang *et al*., 2011, Andrade *et al*., 2013). These investigations have often presented conflicting results (Gianinazzi & Vosátka, 2004, Rowe *et al*., 2007, Faye *et al*., 2013). What these studies highlight is that optimisation of AM inoculation strategies requires specialist management practices. These practices include i) careful inoculum selection to ensure quality and suitability to the host plant, ii) minimal tillage, iii) intelligent crop rotation (non-mycorrhizal should not precede mycorrhizal crops), iv) organic fertilisation practices, and v) avoiding use of agrochemicals i.e. pesticides (Dodd & Ruiz-Lozano, 2012, Berruti *et al*., 2014) wherever possible.

Stricter governance of air pollution has led to greatly reduced inorganic S deposition to soils. A simultaneous increase in high yielding crop varieties has further depleted soil of plant available SO_4_^2-^ stocks (McGrath *et al*., 2003). Soil S exists up to 95% organically bound (Autry & Fitzgerald, 1990), predominantly as aliphatic and aromatic sulfonates (30-70%) and sulfate esters (20-60%), and is not directly available to plants (Zhao *et al*., 2006). Hydrolysis of the sulfate ester bond is facilitated by bacteria and saprotrophic fungi, however, aromatic sulfonate mobilisation is facilitated solely by specific bacteria (Schmalenberger & Kertesz, 2007). Many bacteria isolated from soil possess the ability to mobilise aliphatic sulfonates (King & Quinn, 1997), however, aromatic sulfonate mobilisation has been shown to be more deterministic of plant S nutrition. Indeed, mobilisation of aromatic sulfonates has been shown to promote the growth of tomato plants (Kertesz & Mirleau, 2004) and *Arabidopsis* plants (Kertesz *et al*., 2007). The genetic organisation of this activity is catalysed by a FMNH_2_-dependent monooxygenase enzyme complex encoded in the *ssu* gene cluster (Eichhorn *et al*., 1999). The monooxygenase SsuD cleaves sulfonates to aldehydes and the reduced flavin for this process is provided by the FMN-NADPH reductase SsuE. For aromatic desulfonation an additional *asfRABC* gene cluster is required and the *asfA* gene is a marker for this activity (Vermeij *et al*., 1999, Schmalenberger & Kertesz, 2007).

Studies on aromatic desulfonating bacterial phylogeny identified numerous Beta-Proteobacteria; *Variovorax, Polaromonas, Hydrogenophaga, Cupriavidus, Burkholderia* (now *Paraburkholderia*) and *Acidovorax,* the Actinobacteria; *Rhodococcus* and the Gamma-Proteobacteria; *Pseudomonas* (Schmalenberger & Kertesz, 2007, Schmalenberger *et al*., 2008, Schmalenberger *et al*., 2009, Fox *et al*., 2014). Additionally, *Stenotrophomonas* and *Williamsia* species have been isolated from AM hyphae (Gahan & Schmalenberger, 2015).

Plants rely on microbial populations for mobilisation of otherwise unavailable nutrients from soil. AM fungi have been shown to play an important role in mobilisation of P (Read & Perez-Moreno, 2003) and N (Hodge *et al*., 2001), however, their role in mobilisation of the essential macro-nutrient S remains unclear. AM fungi increase the nutritive absorptive surface area of their host plant via their extraradical hyphae (ERH) providing a niche for interacting with organo-S mobilising bacteria (Barea *et al*., 2002). Additionally, AM symbiosis alters the chemical composition of their host plants exudates which may stimulate functional bacterial populations (Lynch & Whipps, 1990). Addition of methyl-ester sulfonates (MES) to soil was shown to stimulate ERH growth putatively as a result of sulfonate mobilising bacterial metabolites (Vilarino *et al*., 1997). Enhanced ERH growth may stimulate further bacterial proliferation in a potential positive feedback loop.

The hypothesis of this study was that AM inoculation may increase colonisation rates, host plant biomass yield, and alter organo-S mobilising microbial communities. Effectively manipulated, inoculation with AM fungi has potential to enhance output from agricultural land. It is imperative that the land is sustainably managed with limited inorganic fertilisation input whilst utilising the beneficial properties of AM fungi to reduce agrochemical requirement. It is necessary to understand the full potential of AM inoculation practices as demand from agricultural land increases and available stocks of essential macro-nutrients for fertilisation are depleted (Smit *et al*., 2009).

## 2. Materials and Methods

### 2.1. Site description

The soil used to create the subsequent experiments was obtained from Teagasc, Johnstown Castle, Co. Wexford, Ireland (52°16’N, 6°30’W, 30 m above sea level). The soil type is a poorly drained gley soil (pH 6), organic matter (11%), and loamy topsoil (18% clay), classified as Mollic Histic Stagnosol (WRB 2006). The soil used has not received P or S fertiliser since 1968 and has not been ploughed since 1970 (P0-0, site 5A). Swards are mixtures of *L. perenne, D. glomerata,* and various meadow grass species (Tunney *et al*., 2010).

#### 2.1.1. Bi-compartmental Microcosms

Bi-compartmental microcosms were established using 120 g of a one part sand (Glenview Natural Stone, Ireland) to soil (Teagasc, Johnstown Castle) mixture. These systems were established to ascertain the bacterial and fungal community response to inoculation with AM fungi. The bi-compartmental systems were composed of 2 Plexiglas plates (12 x 12 cm) (Access Plastics, Ireland) encasing two 0.5 cm layers of soil separated by a 35 µ m nylon mesh layer (Plastok Associates Ltd, Great Britain). Mesh of this size is sufficiently small to allow mycorrhizal cross colonisation while preventing root access. In the first compartment, *Lolium perenne, Agrostis stolonifera*, or, *Plantago lanceolata* (Emorsgate Seeds, UK) were planted (Figure 1a) and a second root free compartment (Figure 1b) was either; left un-inoculated as a control (C treatment), inoculated with 1.8 g of *Rhizophagus irregularis,* purchased as *Glomus intraradices* (Glomus treatment), or inoculated with a commercial mix of 6 different AM fungi including; *G. intraradices* (now *Rhizophagus irregularis*)*, G. mosseae* (now *Funneliformis mosseae*)*, G. etunicatum, G. hoi, G. constrictum,* and *G. claroidium* (Mixed treatment) (Symbivit, Symbion, Czech Republic).

**Figure 1.**
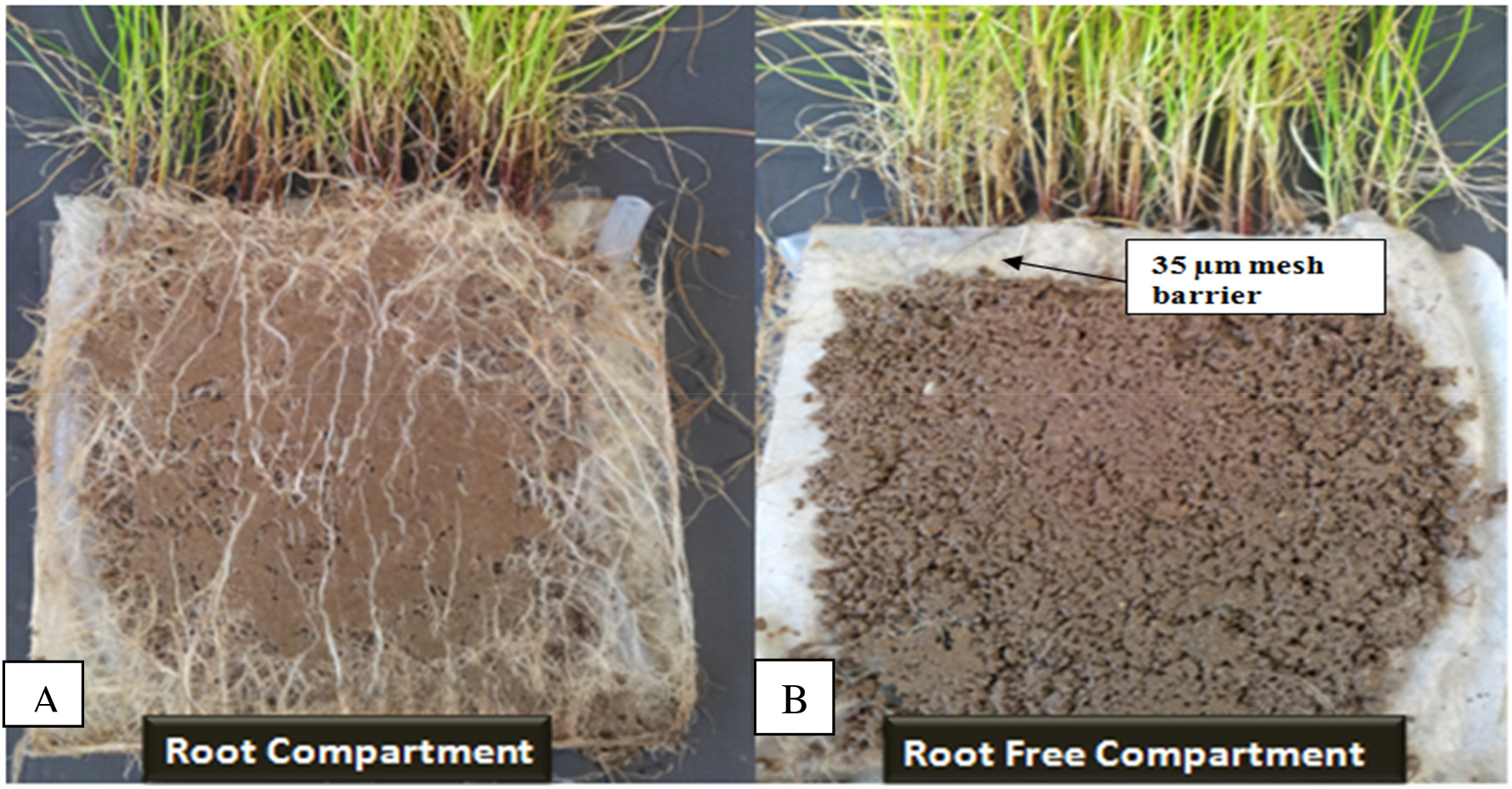
Bi-compartmental microcosms (A) the plant root compartment (B) the root free – arbuscular mycorrhiza compartment, and a selectively penetrable nylon mesh (35 µm).

The commercial AM mixed inoculum is supplied as dry granules containing 150 infective propagules (IP) per g and additives in this inoculum include; natural clay carriers, bio-actives supporting AM symbiosis (chitin, keratin, natural humates, seaweed extract, and ground minerals), and powered biodegradable gel. The Glomus treatment did not contain these additives and the substrate used in this instance was perlite. For all 3 plants, treatments were carried out in quadruplicate and the plant systems were grown in an A1000 Adaptis plant growth chamber (Conviron, Germany) with day-night temperature of 25-15 °C, respectively, a 12 h day length, 70% relative humidity (RH), and 320 µ moles m^-2^s^-2^ photosynthetically active radiation (PAR). The systems were watered and misted with distilled water (dH_2_O) three times a week, and fortnightly supplemented with modified Hoagland’s solution (50% nutrient concentration but without any S, S free) (Hoagland and Snyder 1933). After 6 months of growth, the systems were harvested and analysed for bacterial and fungal community diversity and bacterial sulfonate mobilising activity.

#### 2.1.2. PGP pot experiments

Pot experiments were established using *A. stolonifera* as host plant to determine the impact of AM inoculation practices on PGP (Plant Growth Promotion) and on sulfonate mobilising bacterial communities. Plants of *A. stolonifera* were grown in 200 g of one part sand to soil mixture. Four experiments were established in replicates of 5 (Figure 2).

**Figure 2.**
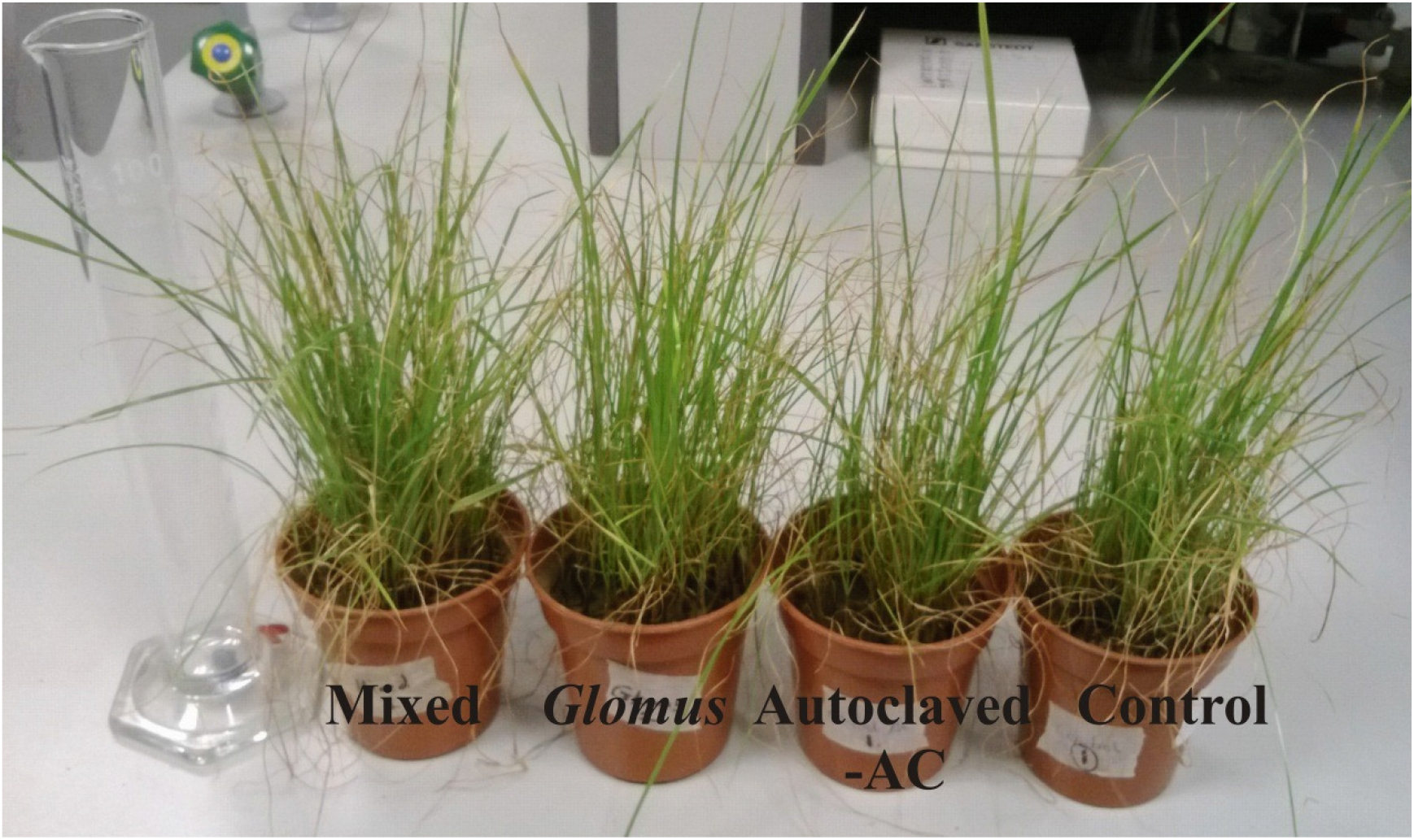
Plant Growth Promotion pot experiments with *Agrostis stolonifera*. Control = free of inoculant, AC = autoclaved inoculum, *Glomus* = *Rhizophagus irregularis* inoculum, Mixed = AM mix inoculum, Bulk Soil = untreated soil.

The first control did not receive AM inoculant to assess the activity of the native AM fungal community and compare that with post-inoculation treatments. The second control received an inactivated commercial mixed inoculum (see above; autoclaved for 15 min, 121 °C) to establish if there was any abiotic element in the inoculum exerting a PGP effect (Autoclaved control = AC). The third treatment received a mono species *Rhizophagus irregularis* inoculation (Glomus), while the final treatment received the viable mixed AM fungi inoculation (Mixed). For all experiments, each pot received 100 *A. stolonifera* seeds and, for all experimental treatments (AC, Glomus and Mixed), 3 g of the respective inoculant. All treatments were grown for 10 weeks in an A1000 Adaptis plant growth chamber (Conviron, Germany) as described above.

### 2.2. Percentage root colonisation

At the time of harvest for both bi-compartmental microcosms and PGP pot experiments, each plant and treatment was examined for percentage root colonisation by AM fungi using a modified version of the grid line intersect technique (McGonigle *et al*., 1990). A representative population of roots (2 root fragments, 10 cm in length were picked from the top, middle, and bottom of the microcosms/pots) and cut into 1 cm segments. The roots were stained with 10% KOH (w/v) for 12 h. The segments were washed with dH_2_O and covered with alkaline H_2_O_2_ to bleach for 60 min. The bleaching solution was discarded and the roots were rinsed thoroughly with water. Roots were acidified in a 0.1 M HCl solution for 12 h to ensure staining of intracellular fungal structures. The HCl solution was discarded and the roots were covered with a lactoglycerol trypan blue stain (lactic acid: glycerol: H_2_O in a 1:1:1 ratio, with 0.05% (w/v) trypan blue) and incubated at 90 °C for 45 min. The stained roots were then removed and covered in lactoglycerol de-stain (minus trypan blue) overnight prior to examination. The 1 cm root segments were examined one field of view at a time (x 1000 magnification). The field of view was moved in reference to a graticule inserted into the eyepiece and the point of intersection was determined at the position of the graticule’s vertical crosshair intersecting the root (McGonigle *et al*., 1990).

Once the point of intersection was noted, the field of view was moved across the surface area of the root and the presence of (1) arbuscules (2) vesicles and (3) hyphae was noted as ‘negative’ (no fungal structures), ‘arbuscules’, ‘vesicles’, or ‘hyphae only’. If the crosshair intersected an arbuscule or vesicle, the respective category was increased by one and the total number of intersections was also increased by one. This was also the case for the ‘hyphae only’ category. In the case of both arbuscules and vesicles being recorded at an intersection, the individual categories were both incremented but the total number of intersections was increased only by one. Arbuscular and vesicular colonisation (AC and VC, respectively) was calculated by dividing their respective counts by the total number of intersections examined i.e. if AC was observed in 50 of 100 intersections: 50/100 = 0.5 or 50%. Hyphal colonisation (HC) was calculated as a proportion of the non-negative intersections. For both bi-compartmental microcosms and pot experiments, 3 replicates of 100 intersections per treatment, per plant were examined to calculate AM fungal colonisation.

### 2.3. Extraction and quantification of bacteria from AM hyphae

#### 2.3.1. Bi-compartmental microcosms

To harvest the bi-compartmental microcosms, in-depth morphological assessment was used to pick 0.5 g of hyphae from the root free compartment of the control experiment and the inoculation treatments using a compound microscope (x 100 magnification) and fine forceps (Hodge & Fitter, 2010). Detailed microscopic observations of the picked hyphae confirmed typical anatomical features of AM fungi and the absence of defined septa (Humphreys *et al*., 2010). As a control, 0.5 g of bulk soil (BS), i.e. soil not under the influence of roots or hyphae, was analysed in parallel. Loosely attached soil was removed from the picked hyphae by dipping in sterile dH_2_O, and associated bacteria were then extracted into 10 mL of sterile saline solution (0.85% w/v) by shaking at 75 rpm on an Elmi Intelli-Mixer RM-2 (Elmi Tech Ltd, Latvia) for 30 min at 4 °C. At this point, an aliquot was taken and used to quantify bacterial communities. Following this, the treatments were centrifuged at 4500 rpm and 4 °C for 20 min and the pellet was frozen immediately (−18 °C) for downstream cultivation independent community analysis.

Using the aliquot of bacterial suspension, 100 µ L was added to 900 µ L of saline solution (0.85% w/v) and a series of 7 tenfold serial dilutions were performed. Most Probable Number (MPN) analysis was undertaken in liquid R2A (Reasoner & Geldreich, 1985) and MM2TS (Fox *et al*., 2014) to enumerate the cultivable heterotrophic and sulfonate mobilising communities. A 20 µ L aliquot from each dilution (10^1^-10^7^) was added to 200 µ L of either R2A or MM2TS in 96 well microtitre plates. Following 2 weeks of growth in an Innova Incubator Shaker Series (New Brunwick Scientific, UK) at 75 rpm and 25 °C, the OD (590 nm) was recorded after 3 minutes shaking (Intensity Level 3) using an ELX808IU spectrophotometer (Bio Tek Instruments Inc., Winooski, VT). The OD_590_ was used to identify the lowest dilution with growth in all five wells, the number of wells with growth in the subsequent two dilutions was used to generate a three digit MPN number and this was used to refer to an MPN table to obtain an MPN g^-1^ value (FDA, 2011). This MPN g^-1^ value was substituted into the following equation: MPN g^-1^ x (1/V) x (DF). V is the volume plated in mL and DF is the dilution factor (0.02 mL and 20, respectively for this experiment).

In order to ascertain the taxonomic diversity of sulfonate mobilisers, a 100 µ L aliquot of bacterial suspension was removed from 3 wells of the liquid MM2TS for the control and each treatment and spread on solid MM2TS under aseptic conditions (Beil et al. 1995) using 6 g L^-1^ of low-S agarose as solidifying agent. After one week, 20 single colonies were selected and sub-cultured for each plant type and treatment. DNA was extracted from each isolate using the quick lysis protocol (Schmalenberger *et al*., 2001) and 16S rRNA gene amplification was undertaken with a final concentration per 25 µ L reaction of 1 X buffer (2 mM MgCl_2_), 0.2 mM dNTP mix, 0.4 µ mol of each primer 27F and 1492R (Lane, 1991) (Table 1) and 0.5 U of DreamTaq polymerase (Fisher Scientific, Waltham, MA). All PCR reactions were carried out in a G-Storm GS2 thermocycler (G-Storm, UK). The cycling conditions were as follows; initial denaturation of 95 °C for 5 min, 32 cycles of 94 °C denaturation (45 s), 55 °C annealing (45 s) and 72 °C extension (90 s). Final extension was undertaken at 72 °C for 5 min.

**Table 1.**
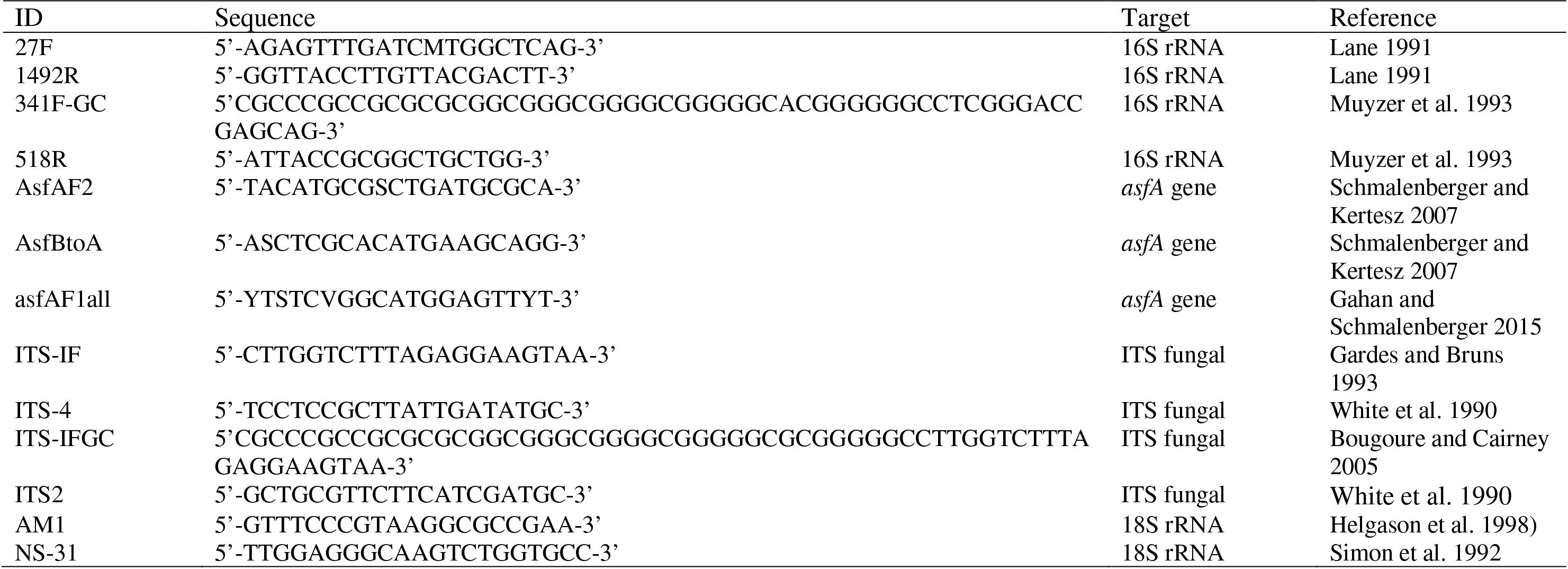
Primers used in this study

To identify dominant OTUs (Operational Taxonomical Units), RFLP (Restriction Fragment Length Polymorphism) was carried out on PCR amplicons using the restriction enzymes *Rsa*I and *Taq*I (5 U per reaction; Thermo Scientific) for 4.5 h at 37 °C. The digested DNA was run on a 2% agarose gel at 100 V for 40 min. Dominating OTUs were screened for presence of the sulfonate mobilising *asfA* gene using the functional *asfA* primers asfAF2 and asfBtoA (Schmalenberger and Kertesz 2007) (Table 1). The *asfA* gene was amplified using the following PCR protocol; initial denaturation of 94 °C for 4 min, 10 cycles of 94 °C denaturation (45 s), 67–57 °C touchdown (30 s), and extension of 72 °C (90 s), plus 25 further cycles at with an annealing temperature of 55 °C. The PCR was conducted using a Kapa 2G Robust PCR kit (Kapa Biosystems, Woburn, MA, USA) in 25 µ L reactions containing; 1 x Buffer A, 1 x Enhancer, 5% DMSO (Sigma-Aldrich), 2 mM MgCl_2,_ 0.2 mM dNTP, 0.5 µ M of each primer and 0.5 U of the Kapa Robust polymerase.

Subsequently, amplified 16S and *asfA* genes of dominant OTUs were purified (GenElute, Sigma-Aldrich, St. Louis, MO) and quantified using a Nano Drop ND-1000 (Thermo Scientific, Waltman, MA, USA) before being subjected to Sanger sequencing (GATC Biotech, Germany) (Sanger and Coulson 1975).

#### 2.3.2. PGP pot experiments

Following a 10-week growth period, the PGP pot experiments were harvested. An in-depth morphological assessment was used to pick 0.5 g of hyphae from the roots of *A. stolonifera* for the control, and the three inoculation treatments (AC, Glomus and Mixed) using a compound microscope (x 100 magnification) (Hodge & Fitter, 2010). Bacteria were extracted and MPN analysis was undertaken in MM2TS media as described above. Additionally, at the time of harvest all above and below ground plant material was dried at 55 °C for 72 h in a fan assisted oven (Carbolite, UK) after which the dry weights were recorded. All excess soil was removed from the roots using a fine bristled brush before drying.

### 2.4. Community fingerprinting

Community fingerprinting analysis was undertaken for the bi-compartmental microcosm experiments. The frozen pellets of the bacterial extractions were used for community DNA extractions using the UltraClean Soil DNA extraction kit from MoBio (Carlsbad, CA). The extracted DNA was used for PCR-DGGE of bacterial 16S rRNA and Fungal ITS communities for the bi-compartmental microcosms.

Bacterial 16S rRNA gene amplification was carried out using the primer pair GC-341F/518R (Table 1) targeting the V3 region for DGGE (Muyzer *et al*., 1993). The final concentration in each 25 µ L reaction was; 1 X buffer (2 mM MgCl_2_), 0.2 mM dNTP mix, 0.4 µ mol of each primer and 0.5 U of DreamTaq polymerase (Fisher Scientific, Waltham, MA). A touchdown PCR protocol was used with the following cycling conditions; initial denaturation of 95 °C for 5 min, 20 cycles of 94 °C denaturation (45 s), 65-55 °C touchdown (45 s), 72 °C extension (45 s), plus 18 further cycles with an annealing temperature of 55 °C. Final extension was undertaken at 72 °C for 5 min. DGGE was carried out on 200 x 200 x 1 mm gels in a TV400 DGGE apparatus (Scie-Plas, Cambridge, UK). Gels of 10% (w/v) acrylamide/bisacrylamide were prepared and run using a linear 30-60% gradient in 1 x TAE buffer (60 °C) for 16.5 h at 63 V (Fox *et al*., 2014). After completion, gels were stained in SYBR Gold (1:10,000 diluted) (Invitrogen, Carlsbad, CA) for 30 min and the image taken on a Syngene G:Box (Cambridge, UK).

Fungal DNA fingerprinting was undertaken using the fungal specific primer ITS-1F (Gardes & Bruns, 1993) and ITS-4 (White *et al*., 1990) (Table 1). This product was tenfold diluted and used as template in a nested PCR using the primer pair ITS-1FGC (Bougoure & Cairney, 2005) and ITS-2 (White *et al*., 1990) (Table 6). Both PCRs were undertaken in 25 µ L reactions each containing; 1 X buffer (2 mM MgCl_2_), 1 M betaine, 0.2 mM dNTP mix, 0.4 µ mol of each primer and 0.5 U of DreamTaq polymerase (Fisher Scientific, Waltham, MA). For both PCR reactions, amplification was performed as follows: initial denaturation of 95 °C for 5 min, 40 cycles at 95 °C denaturation (45 s), 60 °C annealing (45 s) and 72 °C extension (45 s), and 20 further cycles with an annealing temperature of 50 °C (Gardes & Bruns, 1993). Final extension was undertaken at 72 °C for 5 min. DGGE was run as before with a gradient of 35-65%.

### 2.5. Diversity of AM fungi and AsfA

AM and *asfA* gene diversity was analysed by generating a clone library of the Control, Glomus and Mixed treatment of *L. perenne.* For AM fungal diversity, the AM specific primer AM1 (Helgason *et al*., 1998) alongside the universal eukaryotic primer NS31 (Simon *et al*., 1992) (Table 1) were used to target the AM fungal 18S rRNA gene. The final concentration per 25 µ L PCR reaction was; 1 X buffer (2 mM MgCl_2_), 1 M betaine, 0.2 mM dNTP mix, 0.4 µ mol of each primer and 0.5 U of DreamTaq polymerase (Fermentas, Waltman, MA). The PCR was carried out under the following conditions; initial denaturation of 95 °C for 5 min, 30 cycles of 94 °C denaturation (30 s), 58 °C annealing (60 s) and 72 °C extension (90 s). Final extension was undertaken at 72 °C for 5 min.

For diversity of sulfonate mobilising bacteria, the *asfA* gene was amplified with primers asfAF1all (Gahan & Schmalenberger, 2015) and asfBtoA (Schmalenberger & Kertesz, 2007) (Table 1). A touchdown PCR was carried out under the following conditions; initial denaturation of 95 °C for 5 min, 10 cycles of 95 °C denaturation (30 s), 65-55 °C touchdown (30 s) and 68 °C extension (40 s), and 25 further cycles at 55 °C annealing. Final extension was undertaken at 68 °C for 5 min. PCR was formulated in 25 µ L reactions each containing; 1 X Terra PCR Direct Buffer (2 mM MgCl_2_), 0.2 mM dNTP mix, 0.4 µ mol of each primer and 0.7 U of Terra PCR Direct Polymerase (ClonTech Europe, Saint-Germain-en-Laye, France).

The respective PCR products were purified, quantified (see above) and ligated into the cloning vector pJET1.2/blunt (CloneJet, Thermo Scientific). The respective ligation reaction was transformed into *E. coli* DH5α. To ascertain taxonomic diversity of recombinant plasmids containing an insert of the correct size, RFLP was carried out on PCR amplicons using 10U of the restriction enzymes *Alu*I and *Rsa*I (Thermo Scientific) in the provided buffer overnight. Clones with matching restriction patterns were classified as a single genotype (OTU) using Phoretix 1D software (Nonlinear Dynamics, Newcastle upon Tyne, UK). Unique genotypes with more than one representative clone were re-amplified and the purified PCR product was used for sequence identification (GATC Biotech). The sequences obtained were subjected to gene comparison using BLAST (Altschul *et al*., 1990). Sequences of *asfA* were imported into arb (Version 5.2) (Ludwig *et al*., 2004), translated into proteins and integrated into an *asfA* phylogenetic tree (Schmalenberger *et al*., 2010).

### 2.7. Data analysis

Univariate analysis of MPN, root colonisation and PGP data was carried out using SPSS (Version 22.0; IBM, USA). DGGE fingerprinting gels were digitalised and band patterns analysed with the software package Phoretix 1D (Nonlinear Dynamics, UK). Cluster analysis via UPGMA was carried out and obtained band pattern matrices were exported for DCA and permutation tests (Monte-Carlo with 9,999 replicates) as described previously (Schmalenberger *et al*., 2010).

## 3. Results

### 3.1. PGP effect of inoculation with AM fungi

Following a 10 week growth period in pots, there was an increase in the aboveground dry weight of plants inoculated with the Mixed, Glomus and Control-AC over the Control treatment (*P* < 0.05) (Figure 3). Additionally, Mixed treatment had increased aboveground dry weight over the Glomus and Control-AC (*P* < 0.05). There was not a difference between Glomus and Control-AC (*P* > 0.05). The increased aboveground biomass obtained with the Glomus treatment was attributed to the presence of the AM fungus, whereas, the Control-AC inoculum contained commercial additives that may have led to the increased biomass obtained over the control. Mixed treatment had the largest mean aboveground dry weight (1.24 ± 0.05 g) and the Control experiment had the lowest mean aboveground dry weight of (0.90 ± 0.06 g) (Figure 3A).

The belowground dry weight was analysed and increased root/hyphae biomass was observed for Mixed over the Glomus treatment and controls (*P* < 0.05) (Figure 3). There was not a difference in biomass for the Control, Control-AC or Glomus treatments (*P* > 0.05). Mixed treatment had the largest mean belowground dry weight at 2.37 g, while the remaining experiments had a mean dry weight of 1.71 to 1.97 ± 0.18 g (Figure 3B).

**Figure 3A.**
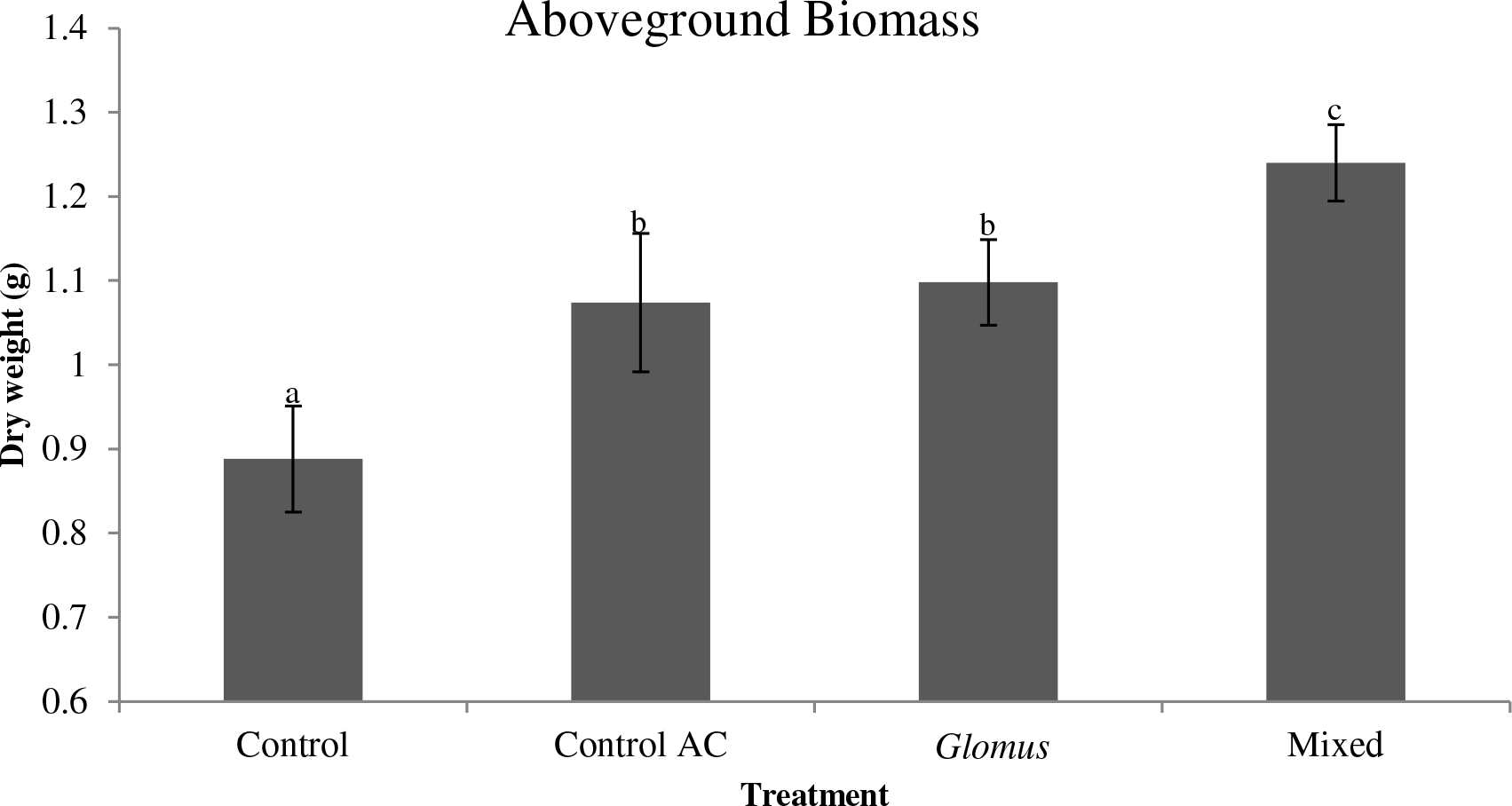
Comparison of the dry weight of *Agrostis stolonifera* aboveground biomass. Control = free of inoculant, Control AC = autoclaved inoculum, *Glomus* = *R. irregularis* inoculum, Mixed = mixed inoculum. Letters (a-c) indicate significant differences.

**Figure 3B.**
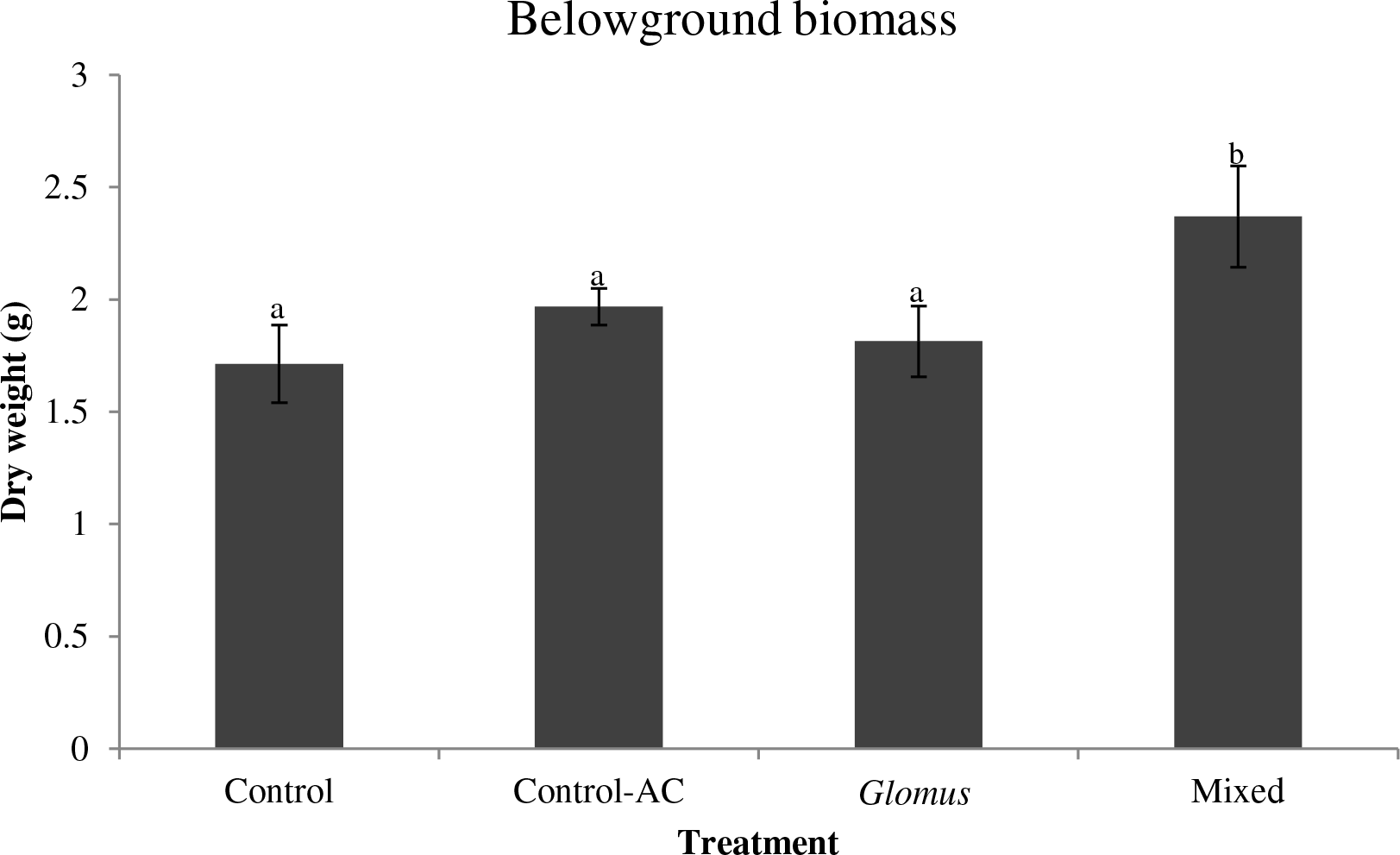
Comparison of the dry weight of *Agrostis stolonifera* belowground biomass. Control = free of inoculant, Control-AC = autoclaved inoculum, *Glomus* = *R. irregularis*inoculum, Mixed = mixed inoculum. Letters (a-b) indicate significant differences.

### 3.2. AM colonisation

#### 3.2.1 Bi-compartmental microcosms

For all three plants, intracellular AC and VC increased following both the *Glomus* and Mixed treatment over the C treatment (*P* < 0.05) (Figure 4). Additionally, there was an increase in AC following Mixed over *Glomus* treatment for *L. perenne* and *A. stolonifera* (*P* < 0.05) (Figure 5A, B), this effect was not observed for *P. lanceolata* (*P* > 0.05) (Figure 4C). There was not an additional increase in VC with Mixed over *Glomus* treatment for the three plants analysed (*P > 0.05).* Finally, there was not an effect of AM inoculation on HC (*P* > 0.05) due to the presence of dense networks of hyphal mycelia surrounding the plant root which made it difficult to identify points of infection (Figure 4).

**Figure 4.**
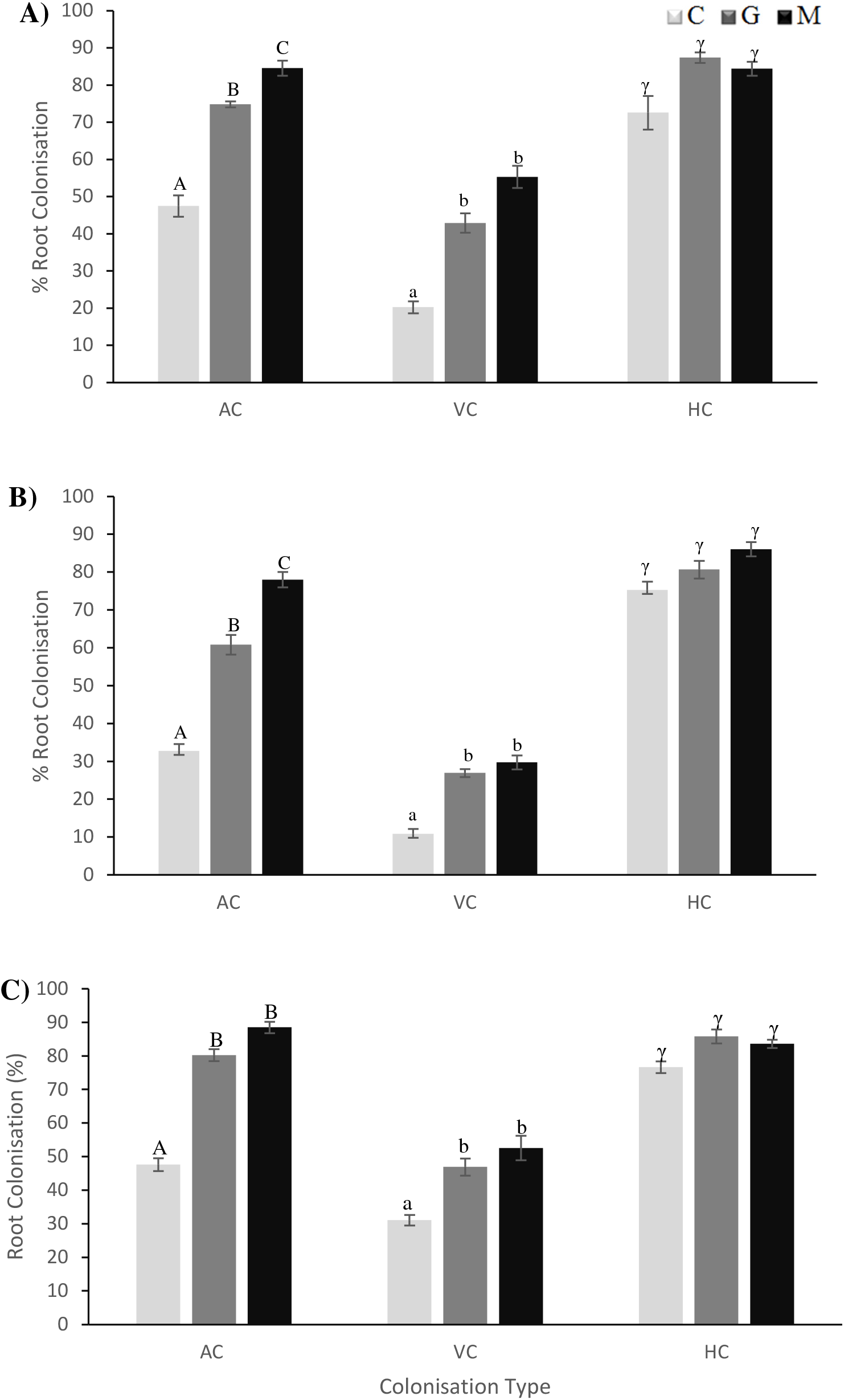
Arbuscular mycorrhizal root colonisation for *Lolium perenne* (A), *Agrostis stolonifera* (B) and *Plantago lanceolata* (C). Arbuscular colonisation = AC, vesicular colonisation = VC, and hyphal colonisation = HC. Treatments include Control = C, Glomus = G (*R. irregularis* inoculum), and Mixed = M (6 AMF species inoculumn). Letters (A-B, a-b, γ) indicate significant differences.

**Figure 5.**
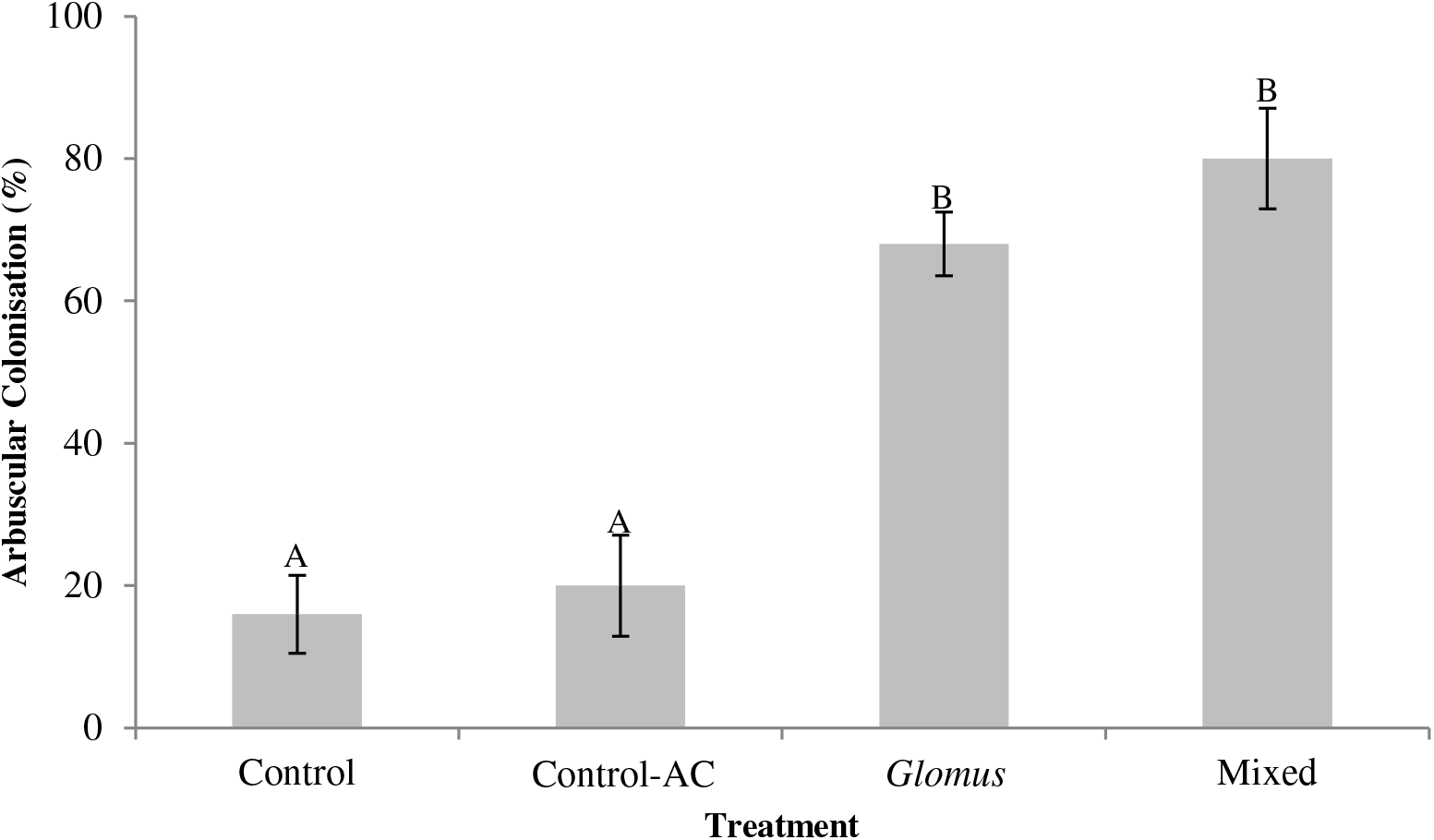
Arbuscular mycorrhizal root colonisation for Plant Growth Promotion (PGP) pot experiments with *Agrostis stolonifera.* Control = free of inoculant, Control-AC = autoclaved inoculum, Glomus *= R. irregularis* inoculum, Mixed = AM mix inoculum. Letters (A-B) indicate significant differences.

#### 3.2.2 PGP pot experiments

The PGP pot experiments demonstrated similar colonisation rates and sensitivity to AM species as was observed with the bi-compartmental microcosms. Arbuscular colonisation was increased following both Glomus and Mixed inoculation over the Control and the Control-AC treatments (*P* < 0.05). There was not an increase in colonisation rate with Mixed over *Glomus* inoculations or Control-AC over the un-inoculated Control (*P* > 0.05) (Figure 5).

Colonisation rates were 16% and 20% for control and Control-AC whereas for Glomus and Mixed treatment the colonisation rate was 68% and 80%, respectively (Figure 5). This result indicates that inoculation with active AM inocula increased colonisation capacity.

### 3.3. Quantification of cultivable heterotrophic microbes

#### 3.3.1 Bi-compartmental microcosms

For all plants, more abundant bacterial communities were observed in R2A and MM2TS media for picked hyphae (Control, Glomus, and Mixed) over untreated bulk soil (*P* < 0.01) (Figure 6). Additionally, higher abundances were observed with Mixed and Glomus treatment over the Control experiment for *L. perenne* and *P. lanceolata,* respectively (*P* < 0.05) (Figure 6A, C). However, an additional increase in abundance of either the total cultivable heterotrophic or the sulfonate mobilising bacterial community was not observed between Mixed and Glomus treatment (*P* > 0.05) (Figure 6A, C). For *A. stolonifera,* there was not an increase in abundance of either the total cultivable heterotrophic or the sulfonate mobilising bacterial community with Mixed or Glomus treatment over the Control experiment (*P* > 0.05) (Figure 6B, C). However, the total cultivable heterotrophic and the sulfonate mobilising bacterial communities were more abundant for picked hyphae and bulk soil for *A. stolonifera* and *P. lanceolata* over *L. perenne* (Figure 6). For *L. perenne,* growth from bulk soil was below the detection limit in MM2TS (1000 MPN g^-1^) (Figure 6A), while for *A. stolonifera* and *P. lanceolata,* it ranged from 10^3^-10^4^ MPN g^-1^ suggesting a higher indigenous community in these plant microcosms (Figure 6B, C).

**Figure 6.**
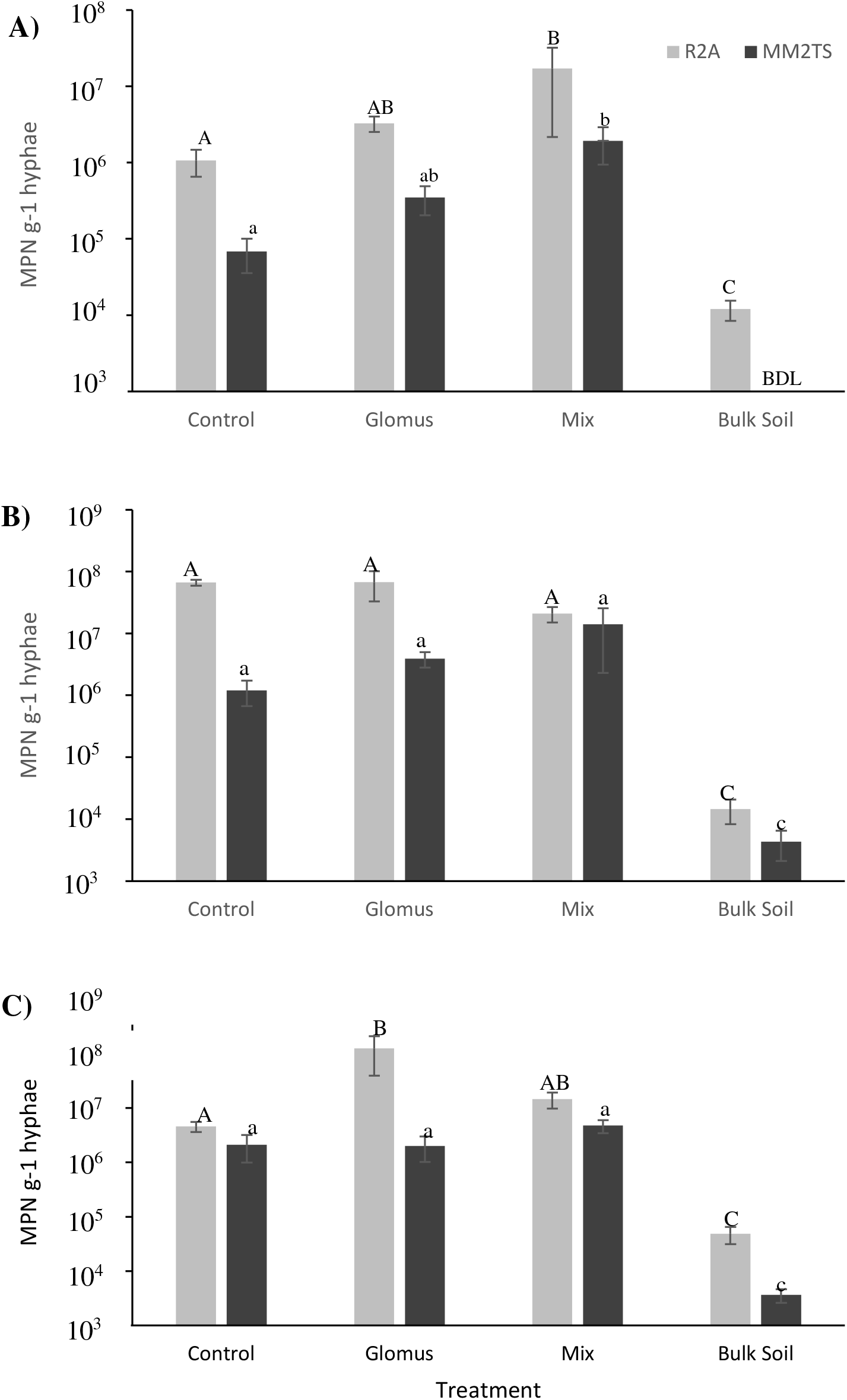
Most probable number (MPN) analysis of bacteria in microcosms with *Lolium perenne* (A), *Agrostis stolonifera* (B) and *Plantago lanceolata* (C). R2A and MM2TS were used to quantify the total cultivable bacterial community and sulfonate mobilisers, respectively. Control = free of inoculant, Glomus = *R. irregularis* inoculum, Mixed = AM mix inoculum, Bulk Soil = untreated soil. Letters (A-B-C, a-b-c) indicate significant differences. Below detection limit = BDL.

#### 3.3.2. PGP pot experiments

For the PGP pot experiment, as observed for the microcosms, the sulfonate mobilising bacterial community abundance was higher for the picked hyphae over the bulk soil (*P* < 0.05) (Figure 7). For the pot experiments unlike the microcosms, a higher abundance of sulfonate mobilisers were observed for both Glomus and Mixed treatments over both the Control and the Control-AC (autoclaved inoculum) treatment (*P* ≤ 0.05) (Figure 7). A difference was not observed between both controls, or between both AMF treatments (*P* > 0.05). Differences between the results obtained for the pot experiments and bi-compartmental microcosms can be attributed to their set-up (pot versus microcosm) and growth period (10 weeks versus 6 months).

**Figure 7.**
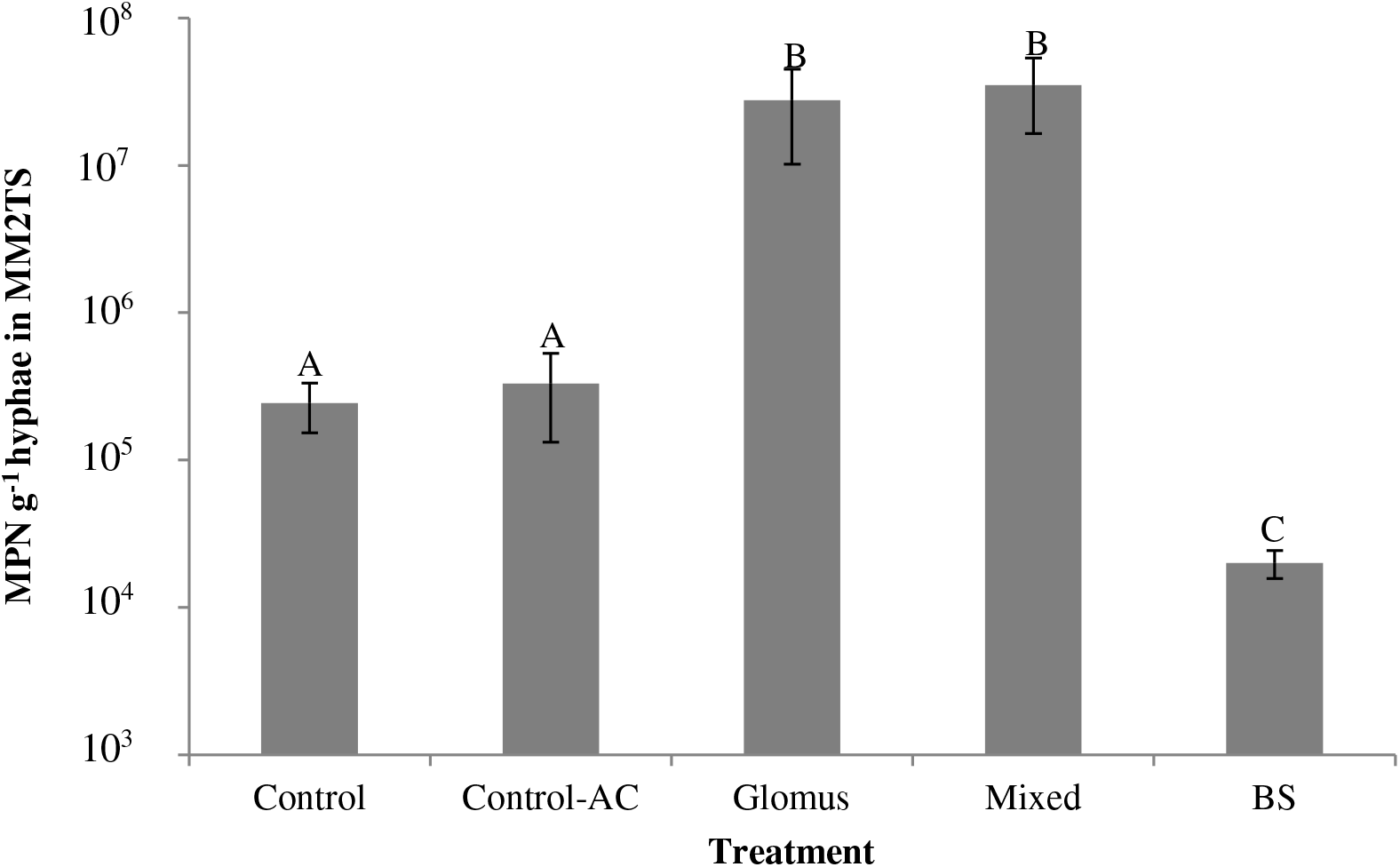
Most probable number (MPN) analysis of the bacterial community in the plant growth promotion pot experiment with *Agrostis stolonifera.* MM2TS were used to quantify the cultivable sulfonate mobilising community. Control = free of inoculant, Control-AC = autoclaved inoculum, Glomus = *R. irregularis* inoculum and Mixed = AM mix inoculum, Bulk Soil = untreated soil. Letters (A-C) indicate significant differences.

### 3.4. Screening of taxonomic diversity of cultivable sulfonate mobilisers

For the 60 isolates taken from the Control, 9 OTUs were obtained in total (based on 16S rRNA gene typing). Of these, 3 were found in *L. perenne,* and 5 in *A. stolonifera* and *P. lanceolata* respectively. Two of these OTUs (OTUs 1 and OTU3) were discovered in all three plants and were present in both the Control and the AMF treatments (Table 2), suggesting that they are common hyphosphere sulfonate mobilisers. The remaining OTUs (OTUs 2, 11, 12, 13, 23, 24, and 25) were present only in the Control experiment and only associated with one of the three plants (Table 2). This result suggests that inoculated soil harbours a distinct community of sulfonate mobilising bacteria.

**Table 2.** Dominant Operational Taxonomic Units (OTUs) of bacteria extracted from the AM hyphae of *Agrostis stolonifera* (*Agrostis*)*, Lolium perenne* (*Lolium*), and *Plantago lanceolata* (*Plantago*) either un-inoculated (Control), inoculated with *R. iregularis* (Glomus), or inoculated with a mix of 6 arbuscular mycorrhizal fungi (Mixed). Symbols mean OTU occurred in either * = all plants, ϯ = all treatments, α = Control, β = Glomus, or γ = Mixed.

| OTU<br>ID | Isolation Source |  |  | Occurrence per Treatment (%) |  |  | Community<br>(%) |
| --- | --- | --- | --- | --- | --- | --- | --- |
|  | *Lolium* | *Agrostis* | *Plantago* | Control | Glomus | Mixed |  |
| 1*† | ✓ | ✓ | ✓ | 55 | 12 | 17 | 25 |
| 2α | ✓ | | | 8 | | | 3 |
| 3*$\alpha\gamma$ | ✓ | ✓ | ✓ | 23 | | 7 | 9 |
| 4*$\beta\gamma$ | ✓ | ✓ | ✓ | | 30 | 10 | 13 |
| 5*$\beta\gamma$ | ✓ | ✓ | ✓ | | 15 | 20 | 12 |
| 6*$\beta\gamma$ | ✓ | ✓ | ✓ | | 23 | 3 | 9 |
| 7β | ✓ | | | | 7 | | 2 |
| 8γ | ✓ | | | | | 12 | 4 |
| 9γ | ✓ | | | | | 7 | 2 |
| 10γ | ✓ | | | | | 3 | 1 |
| 11α | | | ✓ | 3 | | | 1 |
| 12α | | | ✓ | 3 | | | 1 |
| 13α | | | ✓ | 2 | | | 1 |
| 14β | | | ✓ | | | 5 | 1.5 |
| 15$\beta\gamma$ | | ✓ | ✓ | | 5 | 7 | 3.5 |
| 16γ | | | ✓ | | | 2 | 1 |
| 17γ | | | ✓ | | | 2 | 1 |
| 18γ | | | ✓ | | | 2 | 1 |
| 19γ | | | ✓ | | | 2 | 1 |
| 20γ | | | ✓ | | | 2 | 1 |
| 21γ | | | ✓ | | | 2 | 1 |
| 22γ | | | ✓ | | | 2 | 1 |
| 23α | | ✓ | | 5 | | | 1.5 |
| 24α | | ✓ | | 3 | | | 1 |
| 25α | | ✓ | | 3 | | | 1 |
| 26β | | ✓ | | | 5 | | 1.5 |
| 27β | | ✓ | | | 5 | | 1.5 |
| 28γ | | ✓ | | | | 2 | 1 |
| 29γ | | ✓ | | | | 2 | 1 |

For the 60 isolates taken from the Glomus treatment, 8 OTUs were obtained. Of these, 4 were recovered from *L. perenne* (OTUs 4, 5, 6, and 7), 6 from *A. stolonifera* (OTUs 1, 4, 6, 15, 26 and 27), and 5 from *P. lanceolata* (OTUs 1, 4, 5, 6, and 15), respectively. The OTUs 7, 26 and 27 were specific to one of the three plants and the Glomus treatment (Table 2). The remainder of the OTUs overlapped across experiments and plants; however, OTUs 4, 5, 6, and 15 were only present following inoculation in the Glomus or Mixed treatments (Table 2).

For the Mixed treatment, 19 OTUs were obtained. Of these, 6 were obtained from *L. perenne,* and 7 and 12 from *A. stolonifera* and *P. lanceolata,* respectively. The OTUs 1, 3, 4, 5, 6 and 15 were present for more than one plant and more than one experiment (Table 2). There was minimal overlap of OTUs across Control, Glomus and Mixed experiments suggesting that each treatment harbours a distinct cultivable sulfonate mobilising community (Table 2). OTUs 1 and 3 overlapped with the Control experiment, while OTUs 4, 5, 6, and 15 overlapped with the Glomus treatment. The remaining 13 OTUs were exclusive to the Mixed treatment (OTUs 8, 9, 10, 14, 16, 17, 18, 19, 20, 21, 22, 28, and 29). Additionally, the Mixed treatment had a higher number of OTUs than Control and Glomus experiments combined (19 compared to 9 and 8 for Control and Glomus, respectively) suggestive of a higher diversity of sulfonate mobilisers following this inoculation treatment.

### 3.5. Functionality of dominant OTUs

In order to ascertain the sulfonate mobilising functionality of dominant OTUs (see 3.4 above), screening for the presence of the desulfonating *asfA* gene was undertaken. Of 180 bacterial isolates, 29 different restriction patterns were obtained (Table 2), of these, 9 OTUs were cultured from the control (C) experiment, 8 from the Glomus (G) treatment and 19 from mixed (M) treatment. Of the 9 control experiment OTUs, 2 overlapped with Glomus and/or Mixed treatment and 7 OTUs were exclusive to the Control experiment. Of the 8 Glomus treatment OTUs, 5 overlapped with Control and/or Mixed experiment and 3 OTUs were exclusive to the Glomus treatment. Of the 19 Mixed treatment OTUs, 6 overlapped with Control and/or Glomus experiment and 13 OTUs were exclusive to Mixed treatment.

The dominant OTUs were then subjected to sequence identification and classes such as *Alphaproteobacteria, Betaproteobacteria,* and the *Actinobacteria* predominated (Table 3). *Variovorax* was the dominating OTU at 25% of the sulfonate mobilising community and was isolated in the hyphosphere of the Control and the AMF treatments (Table 3). *Paraburkholderia* (13%), *Agrobacterium* (12%), *Polaromonas* (9%), *Rhodococcus* (9%), and *Mesorhizobium* (4%) were also prevalent desulfonating bacteria. Interestingly, while *Polaromonas* (9%) and *Rhodococcus* (9%) have previously been shown to have desulfonating ability, *Paraburkholderia* (13%), *Agrobacterium* (12%), and *Mesorhizobium* (4%) are newly associated with this capability. Furthermore, they were only isolated from Glomus and/or Mixed microcosms (Table 3). All but OTU 6 and 8 (*Rhodococcus* and *Mesorhizobium*) were able to produce an amplification product in the *asfA* PCR (see above).

**Table 3.**
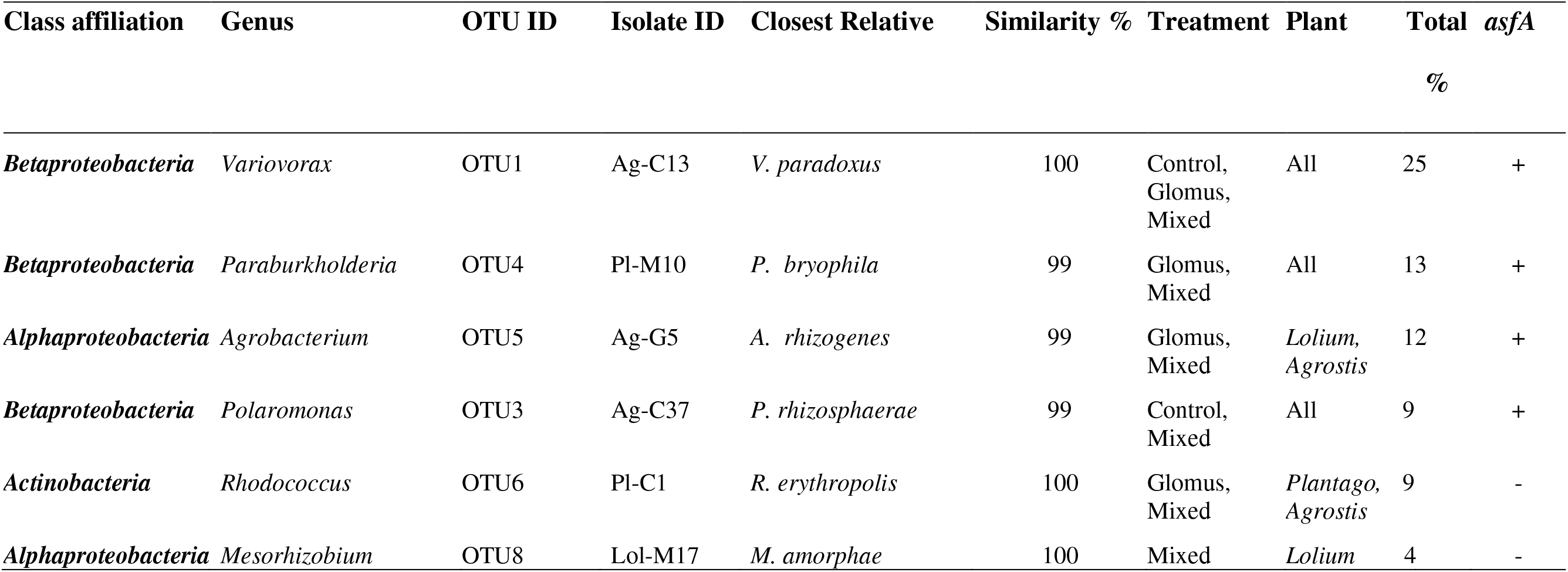
Dominant desulfonating Operational Taxonomic Units (OTUs) obtained from the AM hyphae of *Agrostis stolonifera* (Agrostis), *Lolium perenne* (Lolium), and *Plantago lanceolata* (Plantago) either un-inoculated (Control), inoculated with *R. irregularis* (Glomus), or inoculated with a mix of 6 arbuscular mycorrhizal fungi (Mixed). Isolates were screened for amplification of *asfA* PCR product (+/-)

### 3.6. Community fingerprinting

#### 3.6.1. 16S rRNA community diversity

Community fingerprinting for *L. perenne* (Figure 8A) illustrated that there was a bacterial community shift following both AMF inoculations (Glomus and Mixed) over the Control experiment (*P* < 0.05). Additionally, there was a community shift following Mixed over Glomus treatment (*P* < 0.05). The results show that the bacterial community of *L. perenne* is sensitive to AMF symbiosis and shifts in response to both the Glomus and Mixed treatment.

**Figure 8.**
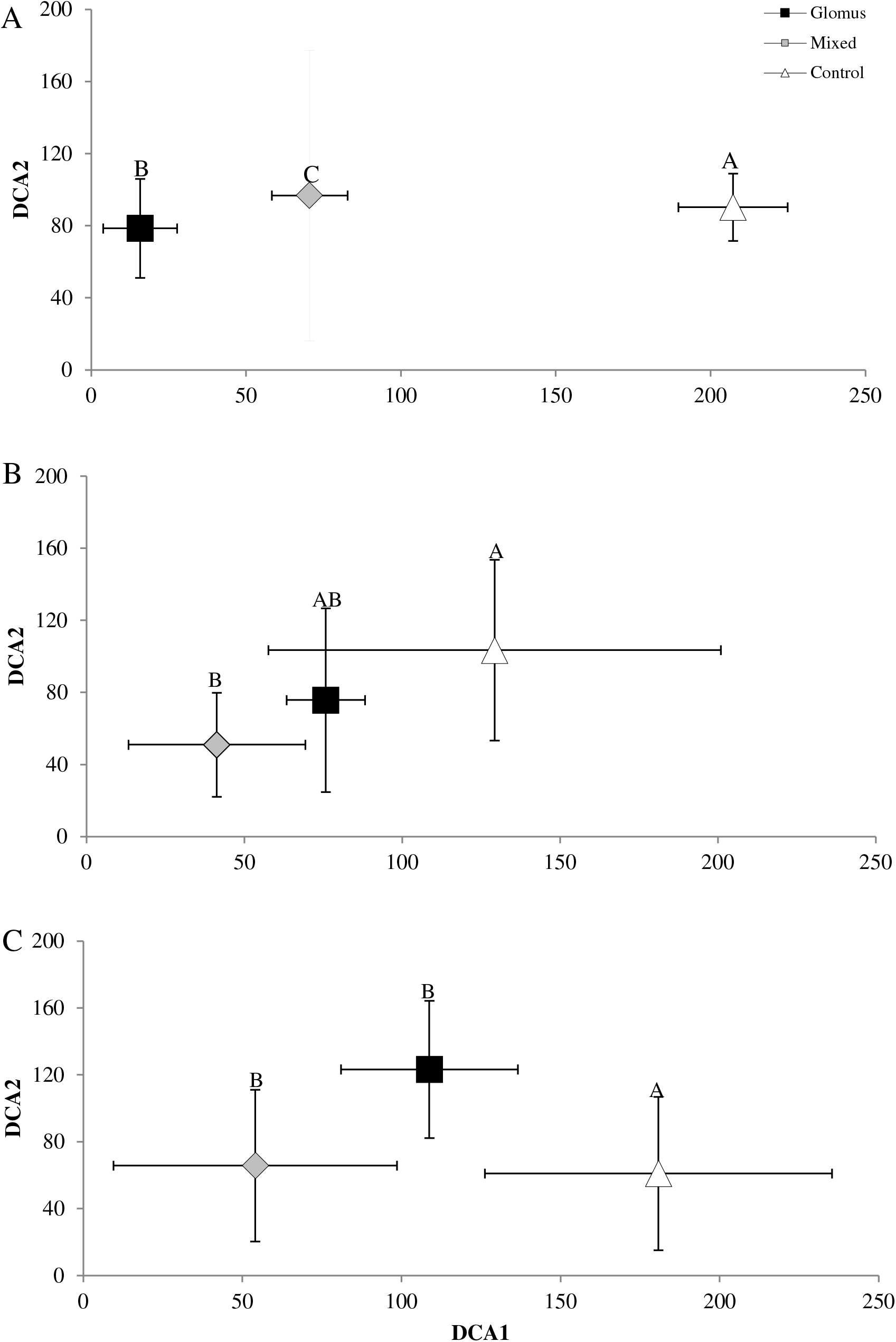
Detrended correspondence analysis (DCA) bi-plot of the 16S bacterial rRNA community fingerprint for *Lolium perenne* (A), *Agrostis stolonifera* (B) and *Plantago lanceolata* (C). DCA1 = Axis 1, DCA2 = Axis 2. Treatments; Control = Triangle, Glomus = Square, Mixed = Diamond. Letters (A-B-C) indicate significant differences.

The bacterial community response of *A. stolonifera* (Figure 8B) illustrated a shift following Mixed over Control experiment (*P* < 0.05). However, there was not a community shift following Glomus over Control or Mixed over Glomus treatment (*P* > 0.05). The results show that the bacterial community of *A. stolonifera* is sensitive to AMF symbiosis; however, this response is dependent on the species of AM fungi used for inoculation.

The bacterial community of *P. lanceolata* (Figure 8C) demonstrated a community shift to both AMF inoculations over the Control experiment (*P* < 0.05). However, there was not an additional shift following Mixed over Glomus treatment (*P* > 0.05). The results obtained for all three plants demonstrate a bacterial community response to AMF inoculation; however, this response is dependent both on the species of AM fungi used to inoculate and the suitability of the plant-AM fungal partnership.

#### 3.6.2. ITS fungal community diversity

Fungal community fingerprinting for *L. perenne* (Figure 9A) illustrated that there was a community shift following inoculation with Glomus treatment over the Control experiment (*P* < 0.05). However, there was not a fungal community shift following Mixed over either Glomus treatment or the Control experiment (*P* > 0.05). Fungal community fingerprinting for *A. stolonifera* and *P. lanceolata* (Figure 9B, C) illustrated that there was a community shift following both AMF treatments over the Control experiment (*P* < 0.05). However, there was not a community shift following Mixed over Glomus treatment (*P* > 0.05).

**Figure 9.**
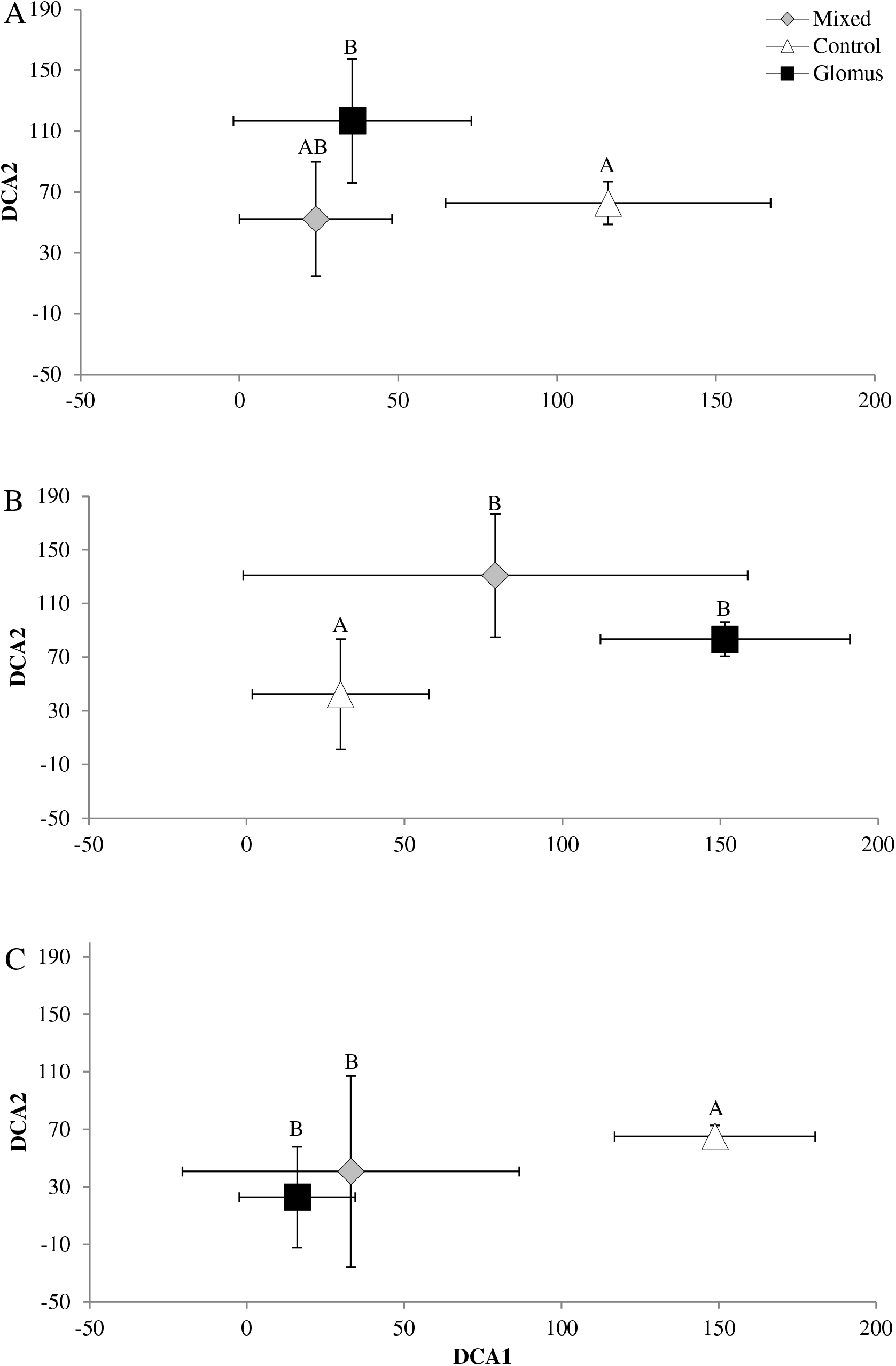
Detrended correspondence analysis (DCA) bi-plot of the fungal ITS region community fingerprint of *Lolium perenne* (A), *Agrostis stolonifera* (B) and *Plantago lanceolata* (C). DCA1 = Axis 1, DCA2 = Axis 2. Treatments; Control = Triangle, Glomus = Square, Mixed = Diamond. Letters (A-B) indicate significant differences.

These results demonstrate that the fungal community in the hyphosphere shifts in response to inoculation with AM fungi but that the response is not necessarily specific to the species of AM fungi present i.e. in the mono species or six species inoculation. As the current fingerprint approach includes both saprotrophic and mycorrhizal fungal species, the presence of AMF bands may give rise to differences in the fungal community composition present for the inoculation (Glomus and Mixed) treatments over the Control experiment.

### 3.7. Diversity of AM fungi and AsfA

#### 3.7.1. Diversity of AM fungi

Clone libraries of AM fungal amplicons from the Glomus and Mixed treatments and the Control experiment of *L. perenne* were screened for different OTUs via RFLP. Screening of 150 clones (50 for each treatment) revealed 6 distinct OTUs with coverage of 100% for each library (Schmalenberger *et al*., 2007). There were no overlapping genotypes/OTUs for the respective libraries. Upon BLAST analysis of dominant OTUs, all were found to be affiliated with members of the Glomeromycota. Thus validating the presence of AM fungi and demonstrating differences across treatments (Table 4). Due to the obligately bi-trophic nature of AM fungi there are limitations in AM sequence availability in reference databases that make taxonomic classification to genus and species level challenging.

**Table 4.** Taxonomic assignment of representative AM fungal Operational Taxonomic Units (OTUs) derived from clonal AM sequence analysis from Control (C), *Glomus* (G) and Mixed (M) treatments of *Lolium perenne*.

| ID | Affiliation | Rep. per<br>OTU | Similarity<br>[%] | Treatment |
| --- | --- | --- | --- | --- |
| C1 | *Acaulospora clone* | 38 | 99 | Control |
| C48 | *Acaulospora clone* | 12 | 99 | Control |
| G1 | *Uncultured mycorrhizal fungus* | 38 | 100 | Glomus |
| G32 | *Acaulospora clone* | 12 | 99 | Glomus |
| M1 | *Acaulospora clone* | 25 | 99 | Mixed |
| M7 | *Scutellospora clone* | 25 | 99 | Mixed |

#### 3.7.2. Diversity of the sulfonate mobilising asfA gene

Clone libraries of the sulfonate mobilising *asfA* gene amplicons from Control, Glomus and Mixed treatments of *L. perenne* were screened for diversity via RFLP. Screening of 150 clones (50 for each treatment) revealed 39 OTUs in total; 17 for the Control (3 overlapping), 18 for Glomus (6 overlapping), and 13 for Mixed treatment (8 overlapping). Library coverage was 94%, 84% and 96% for each respective treatment (Schmalenberger *et al*., 2007). Of these 39 OTUs, 24 OTUs with more than two representatives were subjected to DNA based sequence identification using BLAST, imported into arb (version 5.2), translated into proteins and incorporated into a phylogenetic tree (Figure 10 and 11).

**Figure 10.**
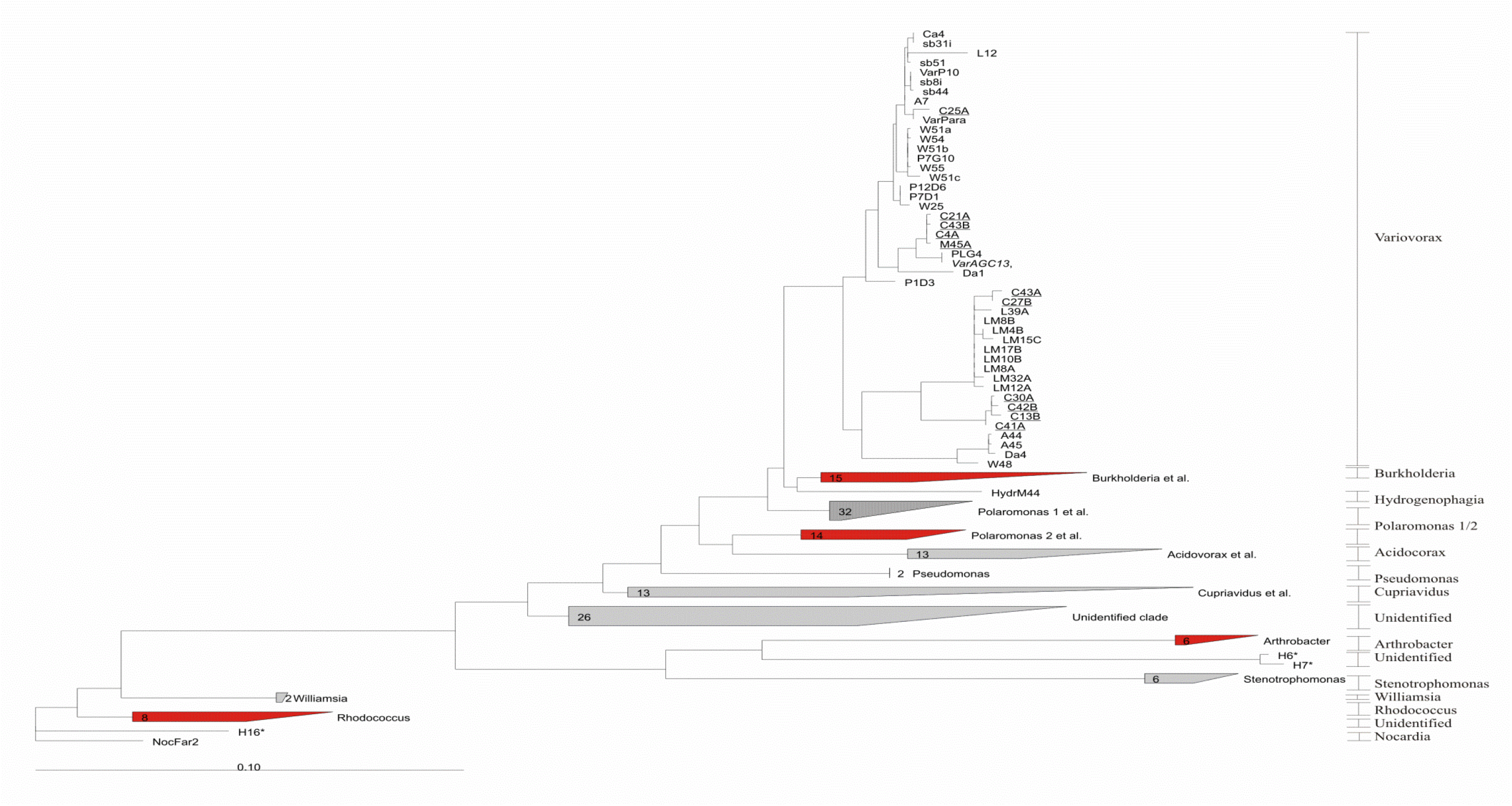
Randomised axelerated maximum likelihood tree of truncated AsfA sequences with the Variovorax clade expanded. C, G and M indicate Control, Glomus and Mixed treatments. Cultivated (*italised*) and molecular isolates from this study are underlined. Molecular isolates from spring barley rhizospheres (sb; (Schmalenberger & Kertesz, 2007)), *Agrostis* grassland rhizospheres (CA; (Schmalenberger *et al*., 2010)), wheat rhizospheres from Broadbalk (W; (Schmalenberger *et al*., 2008)), rhizospheres and soils from the Damma glacier forefield (D; DA; (Schmalenberger *et al*., 2010)) were isolated previously. Highlighted in red are clades to be expanded in Figure 11 below.

**Figure 11.**
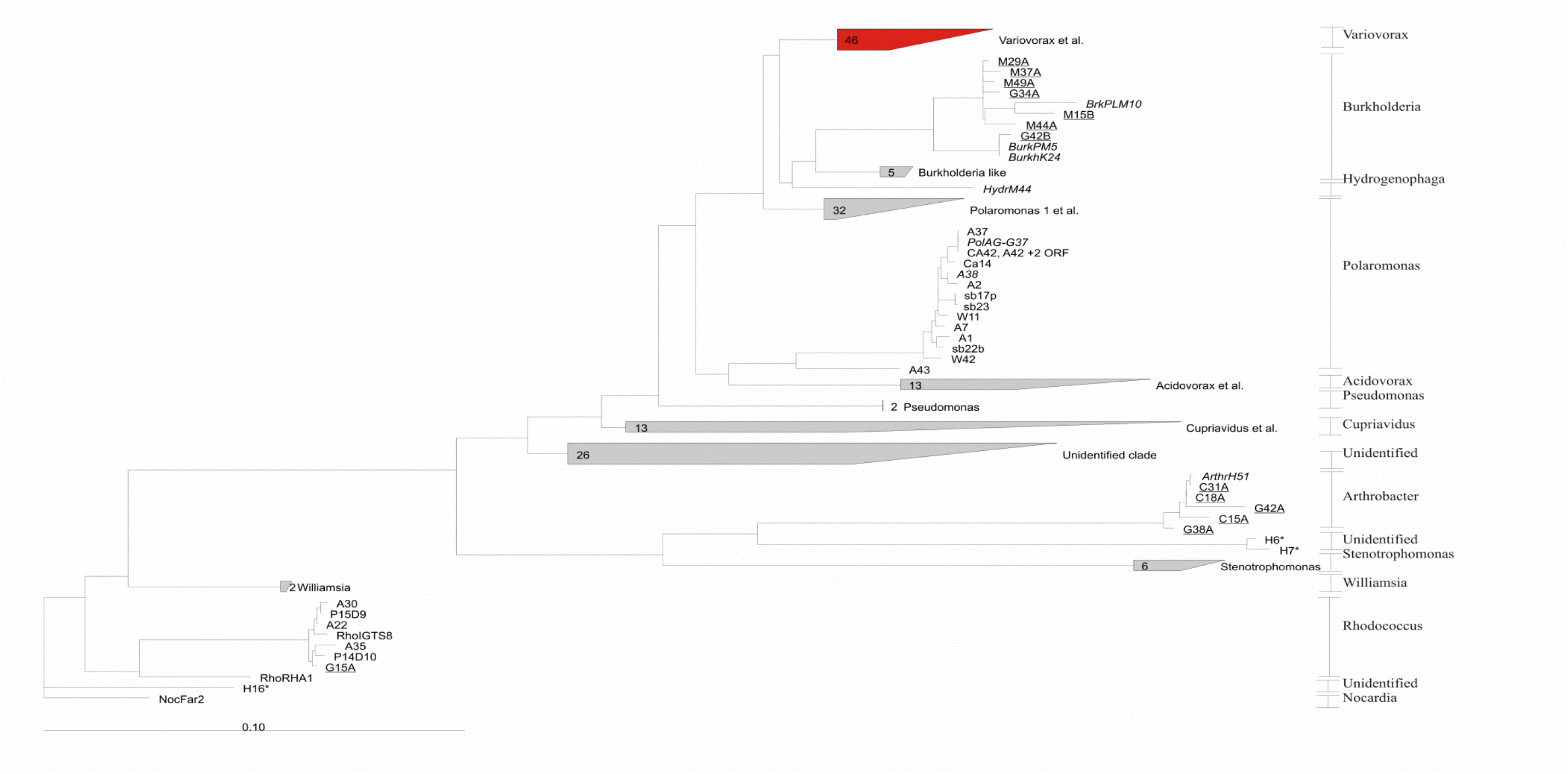
Randomised axelerated maximum likelihood tree of truncated AsfA sequences with the clades minimised in the above figure (red) expanded. C, G and M indicate Control, Glomus and Mixed treatments. Cultivated (*italised*) and molecular isolates from this study are underlined. Molecular isolates from spring barley rhizospheres (sb; (Schmalenberger & Kertesz, 2007)), *Agrostis* grassland rhizospheres (CA; (Schmalenberger *et al*., 2010)), wheat rhizospheres from Broadbalk (W; (Schmalenberger *et al*., 2008)), rhizospheres and soils from the Damma glacier forefield (D; DA; (Schmalenberger & Noll, 2010)) were isolated previously. Highlighted in red are clades expanded in Figure 10 above.

There was a 12% overlap of Control treatment OTUs with Glomus and Mixed treatments and only 3 OTUs occurred in both Control and either AM inoculation treatment (C15A, C18A and C27B) (Table 5). There was a 36% overlap of Glomus and Mixed treatments with 5 overlapping OTUs that dominate the post-inoculation sulfonate mobilising community (G15A, G34A, G42A, M44A, and M45A) (Table 5). The results suggest that the sulfonate mobilising bacterial community composition (*asfA* gene) shifts following inoculation.

**Table 5.**
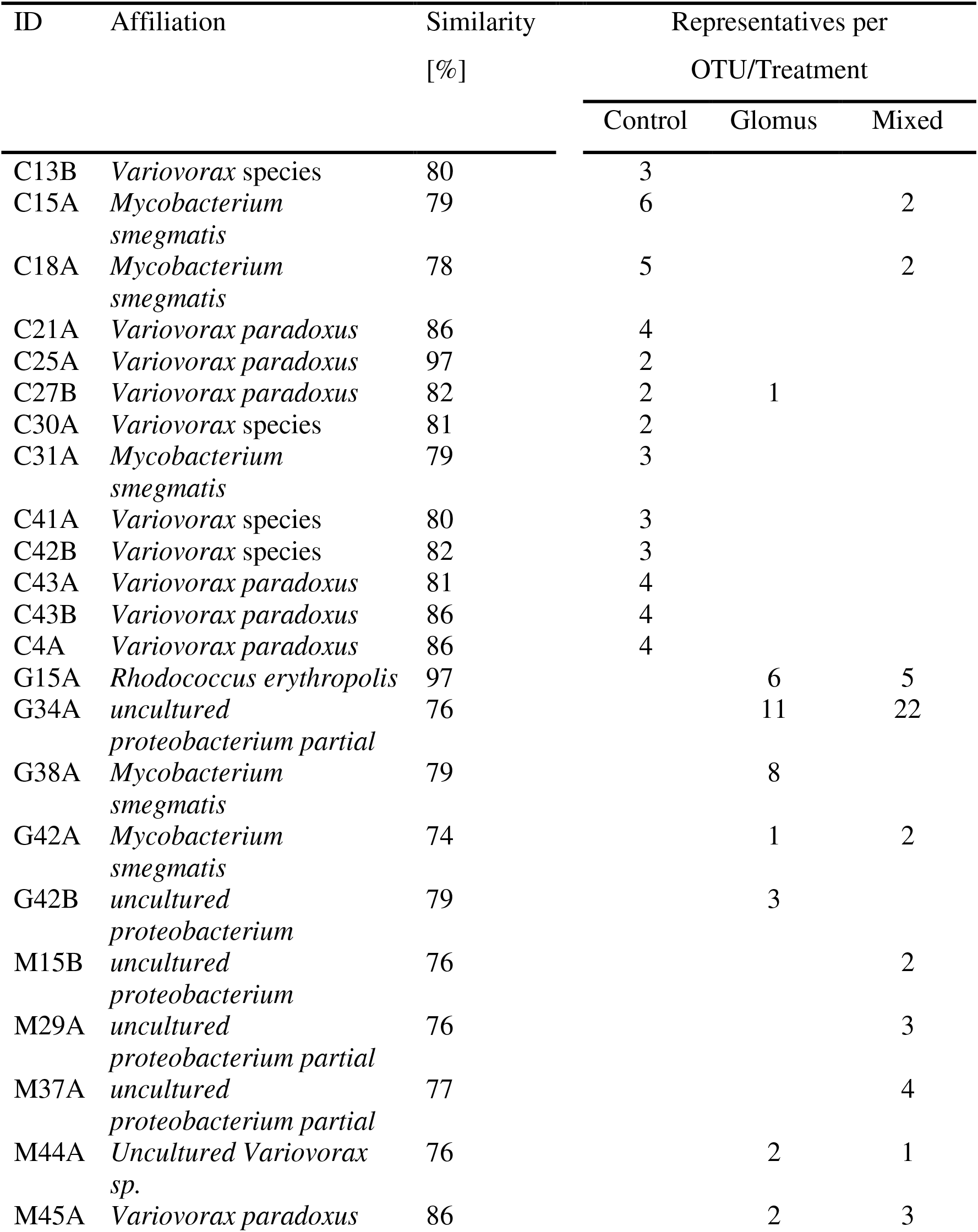
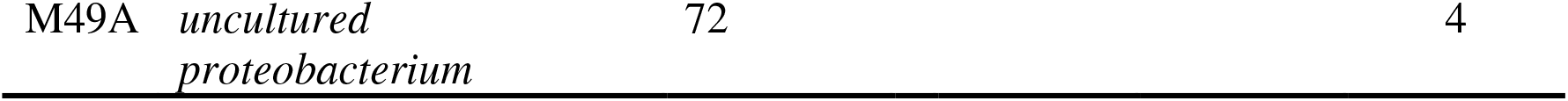
Taxonomic assignment of representative desulfonating OTUs derived from clonal asfA DNA sequence analysis from AM hyphae of *Lolium perenne*, either uninoculated (Control), inoculated with *R. irregularis* (Glomus), or inoculated with a mix of 6 arbuscular mycorrhizal fungi (Mixed).

The sequences obtained were subjected to gene comparison using BLAST (Altschul *et al*., 1990). Sequences of *asfA* were imported into arb, translated into proteins and integrated into an *asfA* phylogenetic tree (Schmalenberger *et al*., 2010) using the randomised axelerated maximum likelihood method. The majority of the Control clones associated closely with the *Variovorax* and *Arthrobacter* clade while the Glomus and Mixed treatment clones were most closely associated to *Paraburkholderia* (for continuity in tree identified as *Burkholderia*)*, Arthrobacter* and *Rhodococcus* clades (Figure 10, 12, respectively). Once again, suggesting that the sulfonate mobilising bacterial community composition is sensitive to inoculation with AM fungi.

## 4. Discussion

Numerous studies have shown that AM inoculation promotes plant growth (Wang *et al*., 2011, Pellegrino *et al*., 2012, Faye *et al*., 2013). Indeed, AM fungi interact synergistically with functional soil microbial populations for P and N mobilisation (Hodge *et al*., 2001, Read & Perez-Moreno, 2003, Hodge & Fitter, 2010). However, rarely has the effect of AM fungal inoculation and AM fungal associated microbiota been examined in the context of organo-S mobilisation. This study aimed to investigate the impact of AM fungal inoculation practices on organo-S transforming microbial community dynamics to potentially improve nutrient supply and promote plant growth. The experiments undertaken demonstrated that AM inoculation significantly increases root colonisation, plant growth, and impacts organo-S mobilising microbial communities.

For the PGP pot experiments, the mono-species *R. irregularis* (Glomus) and mixed AM fungal treatments increased above and belowground plant biomass over the control experiment. Additionally, increases in biomass yield was less pronounced with the mono-species in comparison to the mixed inoculation, this result suggests that mixed AM inoculants have an increased probability of successfully forming an efficiently functional symbiosis. Indeed, many studies corroborate this finding and show that mixed species inoculants exert more beneficial effects in comparison to their mono-species counterparts as a result of increased symbiotic potential arising from multiple species with differential characteristics (Rowe *et al*., 2007, Faye *et al*., 2013, Jin *et al*., 2013). The autoclaved mixed inoculant increased growth more so than the un-inoculated control, however, the PGP effect was significantly less than that observed with the biotic mixed inoculant. Rowe et al. (2007) demonstrated that increased growth can be a result of the substrate in which the inoculant is supplied. Indeed, in an experiment comparing commercial inoculants with 7 different host plants, 5 exhibited increased growth as a result of the inoculant substrate alone thus providing an explanation for the PGP effect obtained with the present AC-Control inoculum (Rowe *et al*., 2007).

AM inoculation significantly increased root colonisation for both the bi-compartmental microcosms and the PGP pot experiments. Tresender (2013) demonstrated a proportionality between increased AM root colonisation and both plant yield and nutrient content (Treseder, 2013), a potential mechanism for which is the increased efficiency of nutrient transfer via characteristic mycorrhizal structures (Van Der Heijden *et al*., 1988, Klironomos & Hart, 2002). An experiment carried out to ascertain the impact of commercial AM fungal inoculant on colonisation rates in *Pisum sativum* demonstrated that mono-species (*G. intraradices*) and mixed AM inoculant (*G. intraradices, G. mosseae,* and *G. clarum*) both significantly increased AM colonisation but there was not an additional benefit of mixed over mono-species inoculation (Jin *et al*., 2013). Despite the equal colonisation capacity of both AMF treatments in the present study, the multi-species (Mixed) treatment increased biomass to a greater extent than the single species (Glomus) treatment. This may be the result of differential nutrient uptake efficiency for multiple distinct AM fungal species in the mixed treatment that is not associated with colonisation capacities and this leading to increased plant biomass yield (Marschner & Dell, 1994).

Cultivable community quantification revealed that total heterotrophic and sulfonate mobilising bacterial communities are larger in the hyphosphere than bulk soil and these findings are consistent with previous studies (Johansson *et al*., 2004, Gahan & Schmalenberger, 2015). For the bi-compartmental microcosms, cultivable heterotrophic and sulfonate mobilising bacterial communities increased in abundance following Mixed treatment for *L. perenne*, only. For *P. lanceolata,* an increased cultivable heterotrophic community was observed in the *R. irregularis* single species treatment (Glomus). There was not an effect of AM inoculation on the sulfonate mobilising community for *P. lanceolata* or *A. stolonifera.* However, the sulfonate mobilising communities were an order of magnitude higher for both plants. The chemical composition of root exudates is dependent on plant species and has been shown to alter microbial community composition and abundance (Lynch & Whipps, 1990, Grayston *et al*., 1998). The characteristic root exudates of *A. stolonifera* and *P. lanceolata* may, therefore, have selectively stimulated larger indigenous communities of sulfonate mobilisers negating the requirement to expend the Control to AM fungi to further stimulate these populations (Jones *et al*., 2004b). For the PGP pot experiment, unlike the bi-compartmental microcosms, increased abundances of sulfonate mobilisers were observed following both Glomus and Mixed inoculations. The duration of this experiment was 10 weeks in comparison to 6 months for the bi-compartmental microcosms. Evidence suggests that S requirement is highest at early stages of vegetative growth (Kertesz *et al*., 2007, Gahan *et al*., 2021), therefore, unlike the bi-compartmental microcosms, the Control may have been allocated to AMF associated microbial populations to facilitate mobilisation of sulfonates to fulfil the plants S requirement.

Bacterial isolates capable of mobilising sulfonates in possession of the *asfA* gene were found to belong to classes such as Alphaproteobacteria, Betaproteobacteria, and the Actinobacteria. Sequences identified include; *Variovorax, Paraburkholderia, Agrobacterium, Polaromonas, Rhodococcus*, and *Mesorhizobium*. *Variovorax, Polaromonas* and *Rhodococcus* species have previously been extracted from the rhizosphere of wheat and have a sulfonate mobilising *asfA* gene (Kertesz *et al*., 2007, Schmalenberger & Kertesz, 2007, Schmalenberger *et al*., 2008, Schmalenberger *et al*., 2009). *Paraburkholderia, Agrobacterium* and *Mesorhizobium* species are newly associated with sulfonate mobilising activity and, furthermore, were only isolated from the AM inoculated treatments due to potential AM specific induced modification of the soil microbial environment (Badri *et al*., 2013). The genera *Paraburkholderia*/*Burkholderia* and *Agrobacterium* have been linked with AM and bacterial interactions and PGP stemming from their ability to mobilise P (Gentili & Jumpponen, 2006). Additionally, *Mesorhizobium* species have been shown to be involved in N fixation (Kaneko *et al*., 2000) and PGP for both chickpea and barley by certain phosphate solubilising strains (Peix *et al*., 2001). Cultivation independent analysis of the sulfonate mobilising *asfA* gene corroborates the cultivation dependent work and revealed minimal overlap of genotypes across treatments. The majority of the Control clones associated closely with the *Variovorax* and *Arthrobacter* clade while the Glomus and Mixed treatment clones were most closely associated to *Paraburkholderia, Arthrobacter* and *Rhodococcus* clades. This result demonstrates that AM inoculation stimulates shifts in the sulfonate mobilising bacterial community. Indeed, AM induced alteration of the sulfonate mobilising bacterial community has been observed previously with DNA extracted from hyphosphere of grassland swards (Gahan & Schmalenberger, 2015).

Community fingerprinting, undertaken for the bi-compartmental microcosms, identified a bacterial and fungal community shift following AMF inoculation. This was expected, as AM fungal symbiosis has been shown to alter microbial community composition in the hyphosphere (Johansson *et al*., 2004, Gahan & Schmalenberger, 2015). Van der Heijden et al., (1998) demonstrated using mono and mixed species inoculations, that growth promotion associated with AMF inoculation was only as good as the most effective AM species in the mix inoculated in isolation. This may provide an explanation for community shifts following the Glomus inoculation but no additional additive shift following Mixed inoculation as observed for *A. stolonifera* and *P. lanceolata*. As observed for root colonisation and cultivation-based analysis of desulfonating communities, *L. perenne* displayed the greatest sensitivity to both AMF inoculation treatments for the cultivation independent community fingerprinting. Helgason et al. (2002) provided an explanation for this by demonstrating that root colonisation, plant-fungal symbiont compatibility and subsequent overall plant vigour were subject to variation depending on the specific AM-plant fungal combination (Helgason *et al*., 2002). It is important, therefore, to select a suitable AMF-plant symbiotic partnership before implementing AMF inoculation practices.

Inoculation with AM fungi can both promote plant growth and stimulate proliferation of bacteria involved in mineralisation of sulfonate-S. Interactions between AM fungi and bacteria have great potential for use in agriculture to improve S supply to plants when available SO_4_^2-^ becomes a limiting factor to growth. The present study also highlights the selective nature of plant receptivity to AM fungi and the importance of careful selection of a high quality AM fungal inoculant suited specifically to the intended ecosystem.

## 6. Acknowledgements

The authors would like to thank the staff of Teagasc, Johnstown Castle, for providing the soil used in this experiment and FP7 People (CIG no. 293429) for funding this project.

## Notes

### Competing Interest Statement

The authors have declared no competing interest.

